# Drought exerts a greater influence than growth temperature on the temperature response of leaf day respiration in wheat (*Triticum aestivum*)

**DOI:** 10.1101/2021.08.16.456456

**Authors:** Liang Fang, Xinyou Yin, Peter E. L. van der Putten, Pierre Martre, Paul C. Struik

## Abstract

We assessed how the temperature response of leaf day respiration (*R*_d_) in wheat responded to contrasting water regimes and growth temperatures. In Experiment 1, well-watered and drought-stressed conditions were imposed on two genotypes; in Experiment 2, the two water regimes combined with high (HT), medium (MT) and low (LT) growth temperatures were imposed on one of the genotypes. *R*_d_ was estimated from simultaneous gas exchange and chlorophyll fluorescence measurements at six leaf temperatures (*T*_leaf_) for each treatment, using the Yin method for non-photorespiratory conditions and the non-rectangular hyperbolic fitting method for photorespiratory conditions. The two genotypes responded similarly to growth and measurement conditions. Estimates of *R*_d_ for non-photorespiratory conditions were generally higher than those for photorespiratory conditions but their responses to *T*_leaf_ were similar. Under well-watered conditions, *R*_d_ and its sensitivity to *T*_leaf_ slightly acclimated to LT but did not acclimate to HT. Temperature sensitivities of *R*_d_ were considerably suppressed by drought, and the suppression varied among growth temperatures. Thus, it is necessary to quantify interactions between drought and growth temperature for reliably modelling *R*_d_ under climate change. Our study also demonstrated that the Kok method, a currently popular method for estimating *R*_d_, underestimated *R*_d_ significantly and should be abandoned.

**Highlight:** Leaf day respiration (*R*_d_) acclimated little to growth temperature, but significantly to drought by reducing its thermal sensitivity. The Kok method underestimates *R*_d_ and should no longer be used.

## Introduction

The ongoing global climate change has resulted in frequent and intense extreme climatic events, such as heat waves, cold snaps and drought spells (Solomon *et al*., 2009; Lloret *et al*., 2012; Perkins-Kirkpatrick & Lewis, 2020; IPCC, 2021). Understanding how these climatic events affect crop physiological processes, particularly photosynthesis and respiration, will be critical for global food security and modelling crop productivity as well as for carbon budgets of agricultural ecosystems in response to climate change (Lobell & Gourdji, 2012; Heskel *et al*., 2013; Yin & Struik, 2017). Respiration plays an essential role in maintaining primary metabolic and physiological functions of plants and costs ca. 40% of gross photosynthetic assimilates of whole plants (Gifford, 1995; Amthor, 2010). Therefore, it strongly affects not only the daily net carbon gain, nutrient acquisition and growth of individual plants, but also the carbon fluxes at ecosystem level (Tcherkez *et al*., 2017a; Tcherkez & Atkin, 2021), in an ever-changing environment.

Respiration occurring in leaves, the metabolically most active plant organs, accounts for a large part of the whole plant respiration (Atkin *et al*., 2007). Leaf respiration is sensitive to short-term (minutes to hours) fluctuations in leaf temperature (*T*_leaf_) and also acclimates to long-term (days) growth temperature changes (Atkin & Tjoelker, 2003). The response of respiration to short-term changes in temperature is often quantified by the parameter called “activation energy” (*E*_a_) of the Arrhenius model or by the *Q*_10_ factor (Atkin & Tjoelker, 2003). The thermal acclimation of leaf respiration has been widely investigated (e.g. Atkin *et al*., 2006; Way *et al*., 2019; Coast *et al*., 2020). The degree to which leaf respiration acclimates to growth temperature differs among species and development stages of leaves (Atkin & Tjoelker, 2003; Atkin *et al*., 2005). Often, leaf respiration acclimates to a sustained warmer growth temperature by decreasing its rate at a reference temperature and/or its thermal sensitivity (i.e. *E*_a_ or *Q*_10_, the slope of the response curve), while acclimation to cooler temperature increases the values of these parameters (Atkin & Tjoelker, 2003). However, this acclimation is not always observed. A recent study that was conducted on the fast-growing species *Eucalyptus globulus* showed an upregulation in basal rates measured at 25°C and *E*_a_ of leaf respiration in warm-grown plants, although the underlying mechanisms are speculative (Crous *et al*., 2017).

Leaf respiration is also modulated by soil water availability, and drought-induced changes in respiration could be associated with changes in availability of substrates (e.g. soluble sugars and carbohydrates), demand for respiratory products (e.g. ATP and NADH) and capacity of respiratory enzymes (Atkin & Macherel, 2009). The impact of drought stress on leaf respiration varies with species, drought severity and drought duration (Flexas *et al*., 2005). In ca. two thirds of the studies reviewed by Atkin & Macherel (2009), leaf respiration was reduced by drought, while in the remaining studies it was unaffected or occasionally increased. Particularly, resilience of leaf respiration in drought was observed under cool and moderate measurement temperatures (Gimeno *et al*., 2010; Gauthier *et al*., 2014). Furthermore, the effect of drought on leaf respiration can also interact with the effects of other environmental factors, for example those of short- or long-term temperature changes and elevated atmospheric CO_2_, especially under field conditions (Ayub *et al*., 2011; Crous *et al*., 2011, 2012; Gauthier *et al*., 2014). These interactions will lead to more unpredictable responses of leaf respiration to drought.

There is growing evidence that the metabolic pathways of leaf respiration vary between illuminated and non-illuminated leaves, as a result of the inhibition of respiration in the light (Tcherkez *et al*., 2017a,b; Tcherkez & Atkin, 2021). Unlike leaf respiration in the dark (*R*_dk_), leaf day respiration (*R*_d_) occurs simultaneously with photosynthetic CO_2_ assimilation and other physiological processes, such as photorespiration, reassimilation and photoinhibition, in the daytime (Yin *et al*., 2020). *R*_d_ is an important parameter in modelling net photosynthetic CO_2_ assimilation (Farquhar *et al*., 1980) and can influence the estimation of other key photosynthetic parameters, such as *V*_cmax_, the maximum rate of Rubisco carboxylation (De Kauwe *et al*., 2016). As the FvCB model (Farquhar *et al*., 1980) is widely used as the basic model for predicting leaf photosynthesis that is to be scaled up to the ecosystem level, *R*_d_ is also crucial in modelling ecosystem gross CO_2_ efflux. However, the different metabolic pathways of *R*_d_ and *R*_dk_ may result in different responses to environmental variables (Gulías *et al*., 2002; Way *et al*., 2019). Compared with the abundance of studies that have explored the environmental impacts on *R*_dk_, the experimental data on how *R*_d_ responds to various environments is generally more lacking, probably because it is difficult to measure *R*_d_.

Techniques to quantify *R*_d_ have been implemented for decades, while there is still debate about the best technique to quantify *R*_d_ (Tcherkez *et al*., 2017a,b; Tcherkez & Atkin, 2021). Either indirect or direct techniques have been developed (Kok, 1948; Laisk, 1977; Haupt-Herting *et al*., 2001; Yin *et al*., 2009; Gong *et al*., 2015; Berghuijs *et al*., 2019). Direct measurement of *R*_d_ (e.g. Haupt-Herting *et al*., 2001; Gong *et al*., 2015) requires sophisticated devices, which are often unavailable. Indirect estimation of *R*_d_ by gas exchange measurements in ecophysiological studies is mostly based on either the Kok method (Kok, 1948) or the Laisk method (Laisk, 1977). The Kok method exploits the Kok effect, that is the abrupt decrease in the slope of photosynthetic CO_2_ assimilation rate (*A*) against irradiance at around the light compensation point (0 - 40 μmol m^-2^ s^-1^). This abrupt switch is interpreted as the consequence of light inhibition to leaf respiration, and thus *R*_d_ can be calculated as the intercept of this linear relationship using points above the break point while the intercept of the linear relationship below the break point is interpreted as *R*_dk_ (Heskel *et al*., 2013; Tcherkez *et al*., 2017b; Yin *et al*., 2020). The Laisk method is based on the FvCB model (Farquhar *et al*., 1980), where the response of *A* to low intercellular CO_2_ concentration (*C*_i_) can be obtained at several (commonly three) levels of irradiances. These curves theoretically intersect at a common point, where the value of *A* represents *R*_d_ and the value of *C*_i_ represents the CO_2_ compensation point in the absence of respiration. The *R*_d_ estimate by the Kok method is often somewhat lower than the estimate by the Laisk method (Villar *et al*., 1994; Yin *et al*., 2011), probably because the Kok method assumes that the PSII electron transport efficiency (*Φ*_2_:) is constant across the light levels used for the gas exchange measurements. A modified Kok method, now known as the Yin method (see Tcherkez *et al*., 2017a) was developed to overcome this weakness of the Kok method, by incorporating the information from chlorophyll fluorescence that accounts for the decline of *Φ*_2_ with light intensity (Yin *et al*., 2009, 2011). The Yin method gave estimates of *R*_d_ that were comparable with those from the Laisk method (Yin *et al*., 2011), with a benefit that the measurements are easier and less time-consuming to implement than those with the Laisk method provided that a fluorometer is available for the concurrent gas exchange and chlorophyll fluorescence measurements.

Although the Kok method, and sometimes also the Yin method, have been applied to common photorespiratory (PR) conditions, theoretically both methods require measurements that are undertaken under nonphotorespiratory (NPR) conditions. This is because both methods using simple linear regression to estimate *R*_d_ implicitly assume that the chloroplast CO_2_ partial pressure (*C*_c_) is constant across light levels whereas a modelling study demonstrated that under PR conditions, *C*_c_ sharply decreased (thus the relative amount of photorespiration increased) with increasing irradiance (Farquhar & Busch, 2017). The significant change of *C*_c_ with irradiance (even if the ambient CO_2_ level is maintained constant) is the result of stomatal and mesophyll regulation of CO_2_ diffusion inside the leaf, ensuring some reassimilation of CO_2_ released by photorespiration and respiration. Using leaf anatomical data combined with 2-D modelling that accounts for CO_2_ diffusion, and thus reassimilation, Berghuijs *et al*. (2019) were able to estimate *R*_d_ based on gas exchange and chlorophyll fluorescence data for PR conditions; they showed that applying the Kok and the Yin methods to PR conditions cause an underestimation of *R*_d_. If the 2-D modelling of the CO_2_ diffusion can reliably estimate *R*_d_, a simpler method, i.e. the coupled FvCB and *g*_m_ (mesophyll conductance) models, can also be explored to estimate *R*_d_ by fitting the coupled model to gas exchange and chlorophyll fluorescence data obtained under PR conditions, because the coupled model can implicitly consider the reassimilation of (photo)respirated CO_2_ (Yin *et al*., 2021). The coupled model has a non-rectangular hyperbolic (NRH) form (von Caemmerer, 2000), and this NRH model was exploited by Yin & Struik (2009) to estimate *g*_m_ from combined gas exchange and chlorophyll fluorescence data, which was shown to be more reliable than the well-known variable J method for estimating *g*_m_. Here, we will test if this model can well estimate *R*_d_ under PR conditions, even if *g*_m_ is unknown beforehand.

Wheat (*Triticum aestivum* L.), as one of the most important staple food crops, is grown worldwide. There is little practical data on how *R*_d_ in C3 Poaceae species like wheat varies with environmental variables, in contrast to the case for tree species, for which studies under either climate-controlled or field conditions have been extensively published in the recent past (e.g. Crous *et al*., 2011, 2012; Way *et al*., 2019; Kumarathunge *et al*., 2020). To date, to our knowledge, no single study provides a set of data of *R*_d_ and its sensitivity to *T*_leaf_ in response to a combination of contrasting water regimes and different growth temperatures in wheat. Understanding how *R*_d_ responds to environmental variables (especially to drought) is critical for quantifying wheat productivity in response to current and future climate change scenarios.

The main objective of this study is to identify the impacts of soil water deficit and growth temperatures on the instantaneous temperature response of *R*_d_ in wheat. We hypothesize that *R*_d_ and its sensitivity to *T*_leaf_ would acclimate to contrasting water regimes and to different growth temperatures, and the thermal acclimation of *R*_d_ may differ between two contrasting water treatments. We estimated *R*_d_ using the established Yin method that requires measurements under NPR conditions. We also aimed to estimate *R*_d_ for common PR conditions; so, the NRH method was explored to estimate *R*_d_ as well. We then evaluated if acclimation differed between PR and NPR conditions. Moreover, we also compared the *R*_d_ estimated by the Kok, Yin and NRH methods.

## Materials and Methods

### Plant materials and growth conditions

To determine any interactive effect of water deficit and genotype on *R*_d_ of wheat plants, an experiment was conducted in a climate-controlled glasshouse at Wageningen University in 2019 (EXP2019). Two genotypes of winter wheat (*Triticum aestivum* L., Thésée and Récital) were used in this experiment. Four batches of seeds were sown with 10-day intervals, creating four replicates. In each replicate, seeds were germinated on a moist filter paper in Petri dishes (one night at room temperature followed by 24 h at 4°C), and then the seedlings were transplanted to multi-cell seedling trays in a glasshouse. When the first leaf fully emerged (ca. 1 week after sowing), plants were moved to a 4°C cold-room (12 h day length, 50 μmol m^-2^ s^-1^ photon flux density) to vernalize for 7 weeks. After vernalization, plants were transplanted to 7-L pots (three plants per pot) filled with a mixture of black soil and peat in a 2:1 (v:v) ratio. The soil mixture was 6 kg per pot and mixed with 1 g N, 1 g P and 1 g K. All potted plants were then moved to a glasshouse and the position of pots were rotated daily and randomly. The climate condition in the glasshouse compartment was set as: 400 ±5 ppm atmospheric CO_2_ concentration, 22/16 ±2°C day/night air temperature (20°C daily average), 75 ±5% relative humidity (corresponding to a vapour pressure deficit (VPD) of 0.66/0.46 kPa for day/night), 16 h photoperiod, and photon flux density at > 400 μmol m^-2^ s^-1^ supplied by sunlight plus supplementary sodium lamps. To avoid any nutrient deficiency, 0.5 g N and 0.25 g N were applied to each pot at the tillering and the stem-elongating stages, respectively.

Another experiment was conducted in a climate-controlled growth chamber in 2020 (EXP2020) to examine whether the leaf day respiration can be altered by different water regimes and growth temperatures. EXP2019 showed no significant difference between the two genotypes in *R*_d_ and its temperature response (see Results); so, only one genotype, Thésée, was used in EXP2020. Considering varying crop durations in different growth temperature regimes, four batches of seeds were sown with 14-day, 17-day and 20-day intervals, respectively, creating four replicates. The same vernalization treatment and plant management practices as in EXP2019 were applied. After vernalization, the potted plants were moved to a climate chamber. The climate settings in the climate chamber were set as: atmospheric CO_2_ concentration, 400 ppm; day/night temperature, 21/17°C; relative humidity, 65%; VPD, 0.87/0.68 kPa for day/night; photon flux density, ca. 410 μmol m^-2^ s^-1^ at soil level; photoperiod, 16 h.

### Experimental treatments

In EXP2019, drought treatment was applied at anthesis in both genotypes. From sowing to anthesis all pots were constantly irrigated to 90% soil water holding capacity, with a gravimetric soil water content of ca. 42%. After anthesis, four replicates of each genotype were well irrigated as control plants (well-watered treatment, WW), whereas the other four replicates of each genotype were subjected to drought stress (treatment DS) by withholding irrigation until the gravimetric soil water content reduced to ca. 16% (drought-stressed treatment, DS). This drought level was maintained until the end of measurements (ca. one week).

In EXP2020, combinations of water deficit treatments and growth temperature were applied. At the booting stage before flag leaves appeared, plants were allocated to three climate chambers with different day/night air temperature settings: high temperature (HT: 28/24°C, average 26.67°C), medium temperature (MT: 21/17°C, average 19.67°C) and low temperature (LT: 16/12°C, average 14.67°C). MT is considered as the control treatment since the temperature was the same as the growth temperature before the temperature treatments started. To minimize any confounding impact of varying VPD, VPD was set identically across chambers (0.87/0.68 kPa for day/night VPD); as a result, 77%, 65%, and 52% relative humidity were applied for HT, MT, and LT treatments, respectively. After anthesis, plants from each climate chamber were subjected to the same two soil water treatments as in EXP2019.

### Gas exchange, chlorophyll fluorescence and leaf N measurements

In both experiments, simultaneous gas exchange and chlorophyll fluorescence measurements were carried out on flag leaves at six leaf temperatures (*T*_leaf_; from 15°C to 40°C with 5°C intervals, except for the LT plants in EXP2020, which were measured at *T*_leaf_ from 12°C to 35°C), 10 days after the onset of the drought treatment, by using a portable photosynthetic system (Li-Cor 6800; Li-Cor Inc., Lincoln, NE, USA) with an integrated fluorescence chamber head of 6 cm^2^. Li-Cor 6800 and plants were together moved to a climate cabinet during measurement to achieve the desired *T*_leaf_. The VPD in the cuvette increased with an increase in *T*_leaf_ and ranged from 1.0 kPa (at 15°C) to 3.0 kPa (at 40°C) for all plants in EXP2019 as well as HT and MT plants in EXP2020. For LT plants in EXP2020, VPD ranged from 1.0 (at 12°C) kPa to 2.5 kPa (at 35°C). For a given *T*_leaf_, incident-irradiance response curves (*A*-*I*_inc_) were assessed under both photorespiratory (PR, i.e. 21% O_2_ combined with 400 ppm ambient CO_2_ (*C*_a_)) and non-photorespiratory (NPR, i.e. 2% O_2_ combined with 1000 ppm *C*_a_) conditions on the same leaf. For the measurements at NPR conditions, a gas cylinder containing a mixture of 2% O_2_ and 98% N_2_ was used. Gas from the cylinder was supplied to the Li-Cor 6800 where CO_2_ was blended with the gas. Photon flux densities in the measurement chamber were 200, 150, 120, 90, 60, 40 and 0 μmol m^-2^ s^-1^ (applied in that order; the value of *A* at 0 μmol m^-2^ s^-1^ of *I*_inc_ represents *R*_dk_) with 5-6 min for each step. The measurements were conducted randomly in each treatment and *T*_leaf_. The operating *Φ*_2_ was determined at each light step as (1-*F*_s_/*F’*_m_) (Genty *et al*., 1989), where *F*_s_ is the steady-state fluorescence, and *F’*_m_ is the maximum fluorescence during the saturating light pulse determined by the multi-phase flash method (Loriaux *et al*., 2013).

After the measurements, the portion of the flag leaves used for gas exchange and chlorophyll fluorescence measurements was cut to measure leaf N elemental content. Rectangle area of the leaf portion was calculated as length multiplied by width which were measured by a vernier caliper. Then the leaf material was weighed after drying in a forced-air oven at 70°C to a constant weight. The concentration of total N in leaf material was analyzed using an EA1108 CHN-O Element Analyzer (Fisons Instruments) based on the micro-Dumas combustion method. From these data, leaf N content on area basis (*N*_area_, g m^-2^) and mass basis (*N*_mass_, mg g^-1^) were calculated.

### Estimation of leaf day respiration

*R*_d_ was estimated by the Yin method (*R*_d(Yin)_) (Yin *et al*., 2009; 2011), where *R*_d(Yin)_ is estimated when *A* is limited by the light-dependent electron transport rate, at which *A* can be described as (Yin *et al*., 2004):

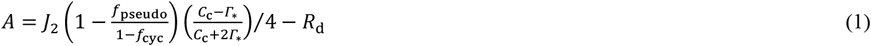

where *J*_2_ is the total rate of e^-^ transport passing PSII, *f*_cyc_ and *f*_pseudo_ represent fractions of the total e^-^ passing PSI that follow cyclic and pseudocyclic pathways respectively, *C*_c_ is the chloroplast CO_2_ partial pressure, and *Γ*_*_ is the *C*_c_-based CO_2_ compensation point in the absence of *R*_d_. By definition, *J*_2_ can be replaced by *ρ*_2_*βI*_inc_*Φ*_2_, where *ρ*_2_ is the proportion of absorbed irradiance partitioned to PSII, *β* is the absorptance by leaf photosynthetic pigments, *I*_inc_ is the incident irradiance, and *Φ*_2_ is the quantum efficiency of PSII electron transport. Then Eqn (1) becomes:

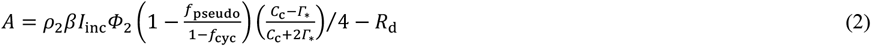

For NPR conditions, *C*_c_ is assumed infinite and/or *Γ*_*_ approaches zero, then Eqn (2) becomes:

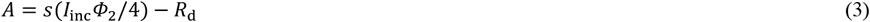

where the lumped parameter calibration factor *s* = *ρ*_2_*β*[1 – *f*_pseudo_/(1 – *f*_Cyc_)]. So, using data of the electron-transport-limited range (200 to 40 μmol m^-2^ s^-1^) under NPR conditions, linear regression plots of *A* against (*I*_inc_*Φ*_2_/4) can be produced, in which *Φ*_2_ is based on chlorophyll fluorescence measurements. The slope of the regression gives the estimate of a calibration factor *s*, and the intercept yields the estimate of *R*_d(Yin)_ under NPR conditions (Yin *et al*., 2009). Obviously, this approach requires that all points are within the linear range of the *A* vs (*I*_inc_*Φ*_2_/4) curves. Here, curves of *A* against (*I*_inc_*Φ*_2_/4) were inspected to exclude the points at high ends that might deviate from the linear pattern, especially for drought plants.

Eqn (2) could also be explored to estimate *R*_d_ under PR conditions if *C*_c_ is maintained constant across irradiance levels. However, it is practically difficult to control *C*_c_ because, before measurements are actually undertaken, one does not know actual values of photosynthetic rate, stomatal conductance and mesophyll conductance required for calculating *C*_c_. Here, the aforementioned non-rectangular hyperbolic (NRH) equation was used to estimate *R*_d_ (*R*_d(NRH)_) for PR conditions. This equation is obtained by combining the well-known FvCB model (Farquhar *et al*., 1980) for e^-^ transport-limited *A* with the Fick’s first law of diffusion for the relation between *A*, intercellular CO_2_ partial pressure (*C*_i_) and *C*_c_:

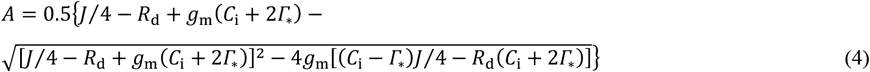

where *g*_m_ is the mesophyll conductance, and *J* is the linear e^-^ transport rate through PSII, which can be calculated as: *J* = *sI*_inc_*Φ*_2_ (Yin *et al*., 2009). The calibration factor *s* was adopted from the slope value of the linear regression for the Yin method from the data under NPR conditions (see above). *Γ*_*_ can be calculated from the relation *Γ*_*_ = 0.5*O*/*S*_c/o_ (where *O* is the concentration of oxygen and *S*_c/o_ is the relative CO_2_/O_2_ specificity factor for Rubisco; Farquhar *et al*., 1980; von Caemmerer, 2013). Nonlinear curve fitting based on Eqn (4) was used to estimate *g*_m_ in a previous study (Yin & Struik, 2009). Here, this NRH equation for *A* was fitted to simultaneously estimate *R*_d(NRH)_ and *g*_m_ under PR conditions. The data used in the NRH method was within the same range of light levels as used in the Yin method.

Use of Eqn (4) combined with *J* = *sI*_inc_*Φ*_2_ to estimate *R*_d_ and *g*_m_ requires that the calibration factor *s* is obtained from strictly NPR conditions. There is no guarantee that this could be the case at extreme conditions (especially when high temperature is combined with drought stress), as the stomatal conductance and *gm* are so low under such conditions that *Γ*_*_/*C*_c_ could not be maintained at the required low level to achieve NPR conditions even when *C*_a_ was set at 1000 ppm. In other words, if a NPR condition cannot be ensured, the obtained *s* is not equal to *ρ*_2_*β*[1–*f*_pseudo_/(1– *f*_cyc_)], but to *ρ*_2_*β*[1–*f*_pseudo_/(1–*f*_cyc_)][(*C*_c_–*Γ*_*_)/(*C*_c_+2*Γ*_*_)]; so, *s* would be lowered by a factor (*C*_c_–*Γ*_*_)*C_c_*+2*Γ*_*_). We introduced dummy variables to Eqn (4) to avoid the confounding effect of measuring under conditions that were not truly NPR and examined whether or not the calibration factor *s* was underestimated. The following identity was verified:

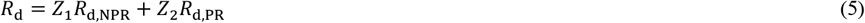

where *R*_d,NPR_ and *R*_d,PR_ are leaf day respiration under NPR and PR conditions, respectively, and *Z*_1_ and *Z*_2_ are dummy variables, which were set in such a way that *Z*_1_ = 1 and *Z*_2_ = 0 correspond to the NPR condition and *Z*_1_ = 0 and *Z*_2_ = 1 correspond to the PR condition. Such a procedure allows to simultaneously estimate the common parameters (*s* and *g*_m_) as well as the different parameters (*R*_d,NPR_ and *R*_d,PR_) from fitting to the combined data obtained under both NPR and PR conditions. This procedure for estimating the common parameters *s* and *g*_m_ is equivalent to the method of simultaneous fitting of the calibration factor and *g*_m_ (e.g. Pons *et al*., 2009) if one is not sure whether a NPR state is reached.

As stated earlier, like the Laisk method, the Kok method is a popular method to estimate *R*_d_ indirectly (e.g. still used recently by Way *et al*., 2019). As our data also allow an implementation of the Kok method, here we compare the NRH, Yin and Kok methods in estimating *R*_d_. As the Kok method has been applied to estimate *R*_d_ (*R*_d(Kok)_) under both PR and NPR conditions (e.g. Tcherkez *et al*., 2017a), for the comparative purpose we also apply the Yin method to the PR conditions by fitting a linear regression to data at the same range of irradiances. It is worthy to note that Eqn (4), upon which the NRH method is based to estimate *R*_d_, can only be applied to the PR conditions.

### The temperature responses of parameters

The thermal response of *R*_dk_ and *R*_d_ was described by the Arrhenius equation normalized with respect to its value at 25°C:

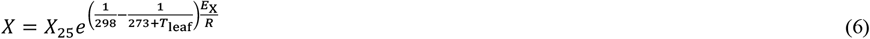

where *X*_25_ represents the value of parameters estimated at 25°C (*R*_d25_ and *S*_c/o25_), *E*_X_ is the activation energy of relevant parameters to temperature (*E*_Rd_ and *E*_Sc/o_; in kJ mol^-1^), and *R* is the universal gas constant (0.008314 kJ K^-1^ mol^-1^). *S*_c/o_ and its temperature response are generally considered to be conserved among C3 species (von Caemmerer *et al*., 2009), so we adopted the value of Cousins *et al*. (2010) for *S*_c/o25_ (3.022 mbar μbar^-1^), and the value of Bernacchi *et al*. (2002) for *E*_Sc/o_, which mathematically equals to the negative of the activation energy for *Γ*_*_: 24.46 kJ mol^-1^. Then the value of *S*_c/o_ at each *T*_leaf_ could be estimated. Sensitivity analysis showed that unlike *g*_m_, the estimated *R*_d_ and its temperature response varied little with variations of *S*_c/o25_ and *E*_Sc/o_ within the physiologically relevant ranges.

### Model analyses and statistics

Simple linear regressions in the Yin method were performed using Microsoft Excel. Non-linear curve fitting procedures in the NRH method and the Arrhenius equation were carried out using the GAUSS method in PROC NLIN of SAS (SAS Institute Inc, Cary, NC, USA). The SAS codes can be obtained upon request to the corresponding author.

## Results

The approach combining the NRH method and dummy variables was used to examine if there was a confounding effect of any under-estimation of the calibration factor *s* on *R*_d(NRH)_ for PR conditions. Using methods either with or without dummy variables, we found that overfitting occurred for a few *T*_leaf_ values of drought plants due to the variability of data, which resulted in the failure of estimating *g*_m_. For such cases, nevertheless, the estimations of *s* and *R*_d(NRH)_ were still reasonable. The results of each parameter (calibration factor *s*, *g*_m_, *R*_d(NRH)_ for PR and *R*_d(Yin)_ for NPR conditions) from approaches with and without dummy variables were very similar (Fig. S1). On average, *s* was underestimated only by ca. 1% (Fig. S1b), suggesting that the gas mixture we used (2% O_2_ combined with 1000 ppm *C*_a_) for estimating the calibration factor did allow to reach a nearly non-photorespiratory state, even for extreme conditions (drought combined with high temperatures). So, for the sake of simplicity, we only present and discuss the results obtained from the method without using the dummy variables.

### Comparisons of R_d_ and its inhibition by light estimated by different methods

*R*_d(Kok)_ was generally lower than *R*_d(Yin)_ under both NPR (18.2% lower) and PR (13.9% lower) conditions (Fig. 1A,B). Under PR conditions, values of *R*_d(Kok)_ were only 66.5% of *R*_d(NRH)_, and those of *R*_d(Yin)_ were 79.6% of *R*_d(NRH)_ (Fig. 1C,D). These findings suggest that the Kok method underestimated *R*_d_ substantially, and the Yin method, if also applied to PR conditions, also underestimated *R*_d_, but less so than the Kok method. Therefore, the light inhibition of leaf respiration, estimated as (1–*R*_d_/*R*_dk_), depended on the methods used for estimating *R*_d_. Under NPR conditions, the estimated average light inhibition by the Kok method was 18.7%, whereas that by the Yin method was only 1.4% (Fig. 2A,B). Under PR conditions, the estimated average light inhibition by the Kok, Yin and NRH methods was 40.3%, 28.5% and 10.1%, respectively (Fig. 2C,D,E), indicating that light inhibition was stronger under PR than NPR conditions. In general, values of *R*_dk_ and *R*_d_ under NPR conditions were greater than under PR conditions (Fig. 3), which was in agreement with results from previous report where both *R*_dk_ and *R*_d_ were higher at 2% O_2_ than at 21% O_2_ in mature leaves (Buckley *et al*., 2017).

**Figure 1.**
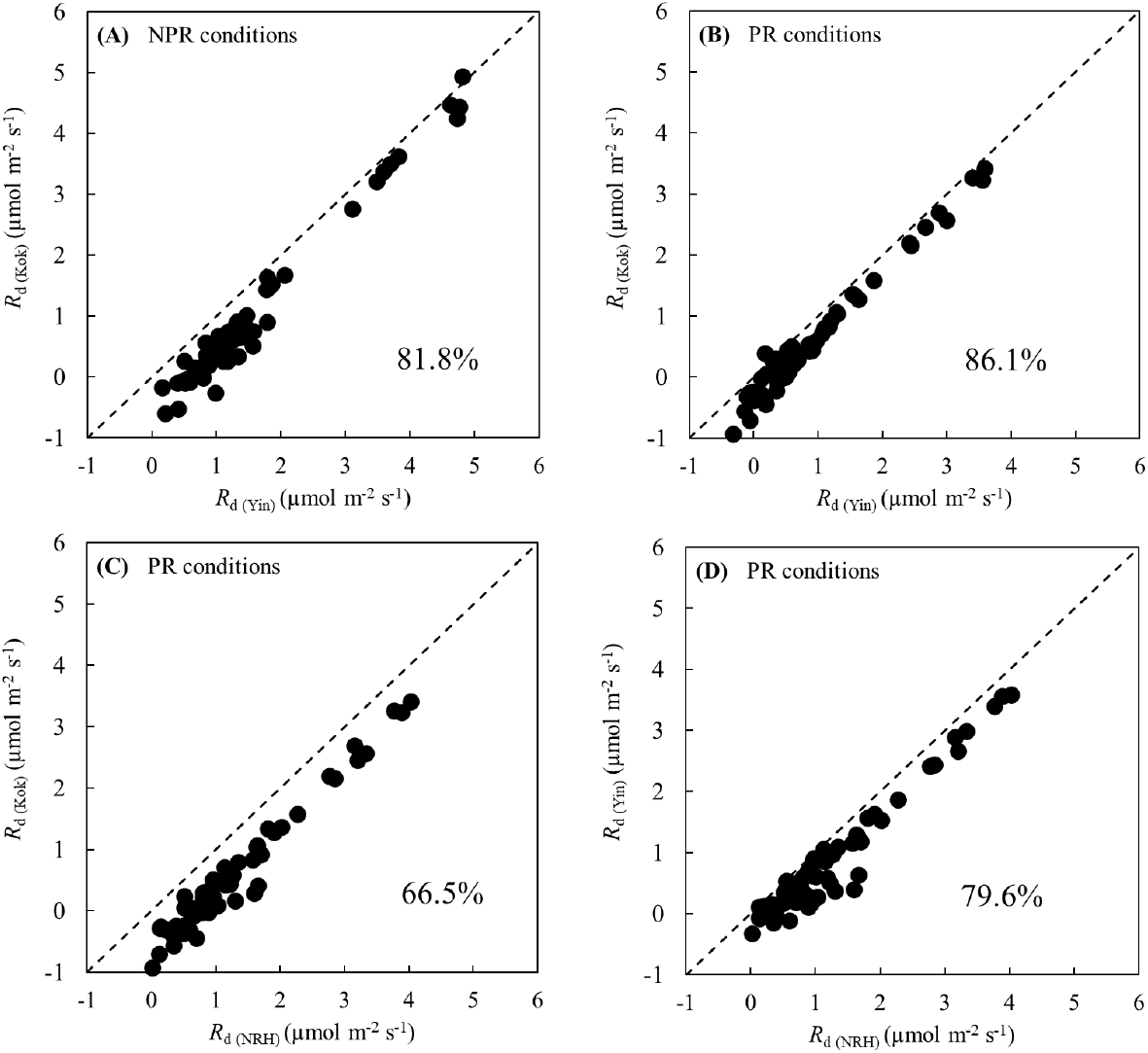
Correlations between leaf day respiration estimated by the Kok method (*R*_d(Kok)_) and estimated by the Yin method (*R*_d(Yin)_) under non-photorespiratory (NPR; panel A) or photorespiratory (PR; panel B) conditions, and correlations between *R*_d(Kok)_ (panel C) or *R*_d(Yin)_ (panel D) and leaf day respiration estimated by the NRH method (*R*_d(NRH)_) under PR conditions, across wheat genotypes Thésée and Récital, growth temperature and water treatments, and leaf temperatures during measurements. Note that the Yin method suits to estimate *R*_d_ for the NPR conditions only; as stated in the text, it is applied also to PR conditions here merely for the comparison purpose. The dashed diagonal represents the 1:1 relationship. The number (%) in each panel is the average of y-axis relative to *x*-axis values.

**Figure 2.**
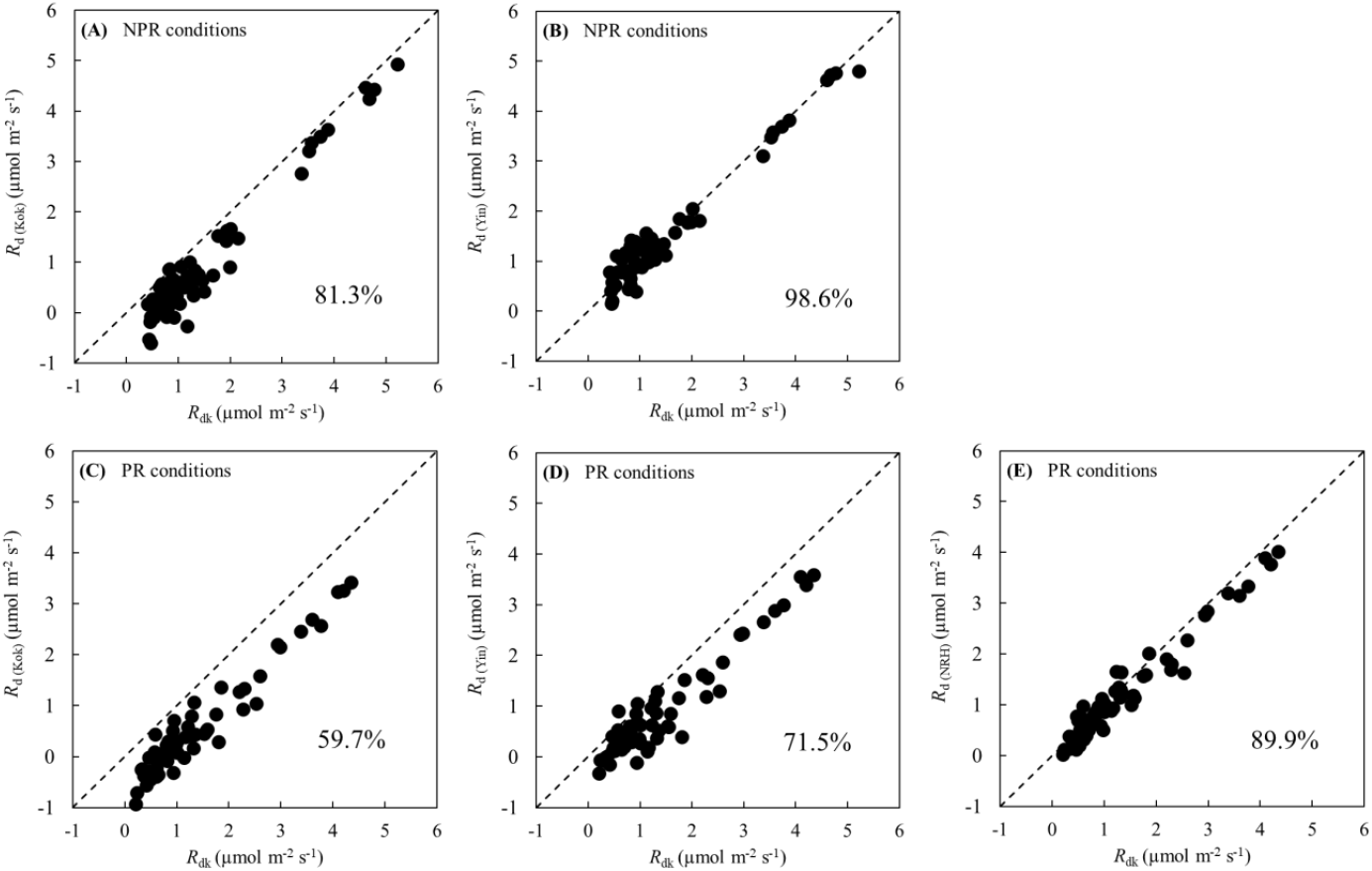
Correlations between leaf day respiration estimated by the Kok (*R*_d(Kok)_), Yin (*R*_d(Yin)_) or NRH (*R*_d(NRH)_) methods and leaf respiration in the dark (*R*_dk_) under non-photorespiratory (NPR; upper panels) and photorespiratory (PR; lower panels) conditions, across wheat genotypes Thésée and Récital, growth temperature and water treatments, and leaf temperatures during measurements. Note that the Yin method suits to estimate *R*_d_ for the NPR conditions only; as stated in the text, it is applied also to PR conditions here merely for the comparison purpose. The dashed diagonal represents the 1:1 relationship. The number (%) in each panel is the average of y-axis relative to *x*-axis values.

**Figure 3.**
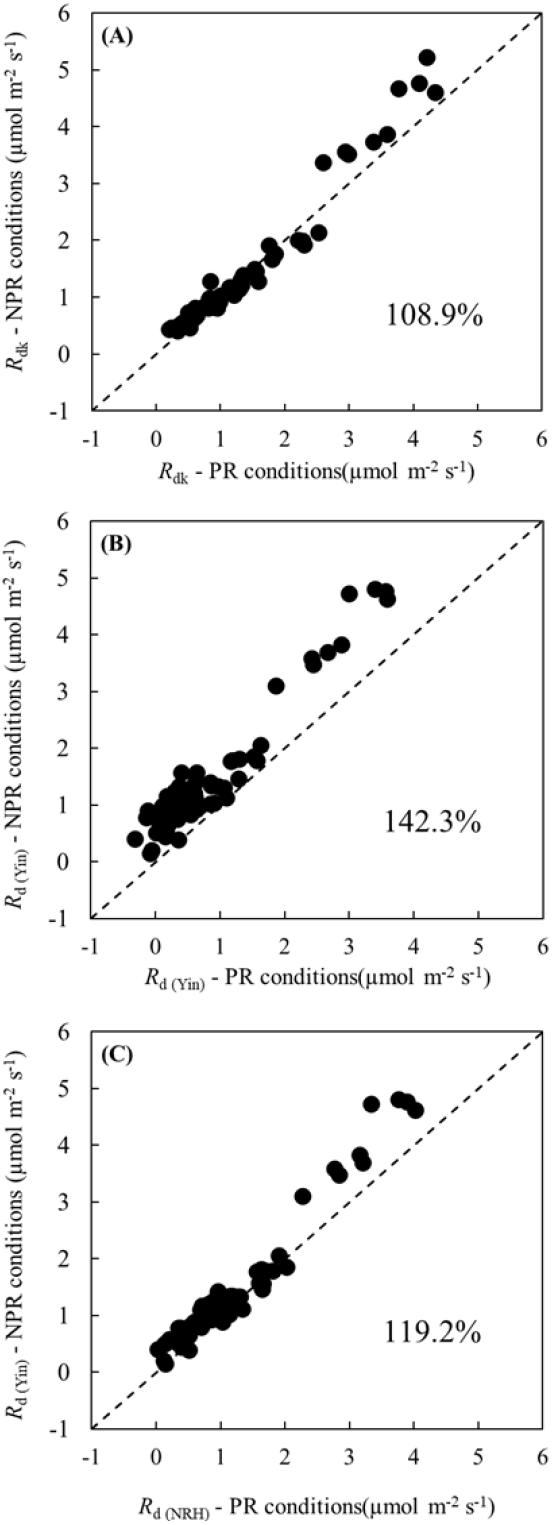
Correlations between leaf respiration in the dark (*R*_dk_) under NPR conditions versus *R*_dk_ under PR conditions (panel A), and correlations between leaf day respiration estimated by the Yin method (*R*_d(Yin)_) under NPR conditions versus *R*_d(Yin)_ under PR conditions (panel B) or that estimated by the NRH method (*R*_d(NRH)_) under PR conditions (panel C), across wheat genotypes Thésée and Récital, growth temperature and water treatments, and leaf temperatures during measurements. Note that the Yin method suits to estimate *R*_d_ for the NPR conditions only; as stated in the text, it is applied also to PR conditions here merely for the comparison purpose. The dashed diagonal represents the 1:1 relationship. The number (%) in each panel is the average of y-axis relative to *x*-axis values.

The obtained thermal responses of *R*_dk_, *R*_d(Kok)_ and *R*_d(Yin)_ under PR and NPR conditions as well as *R*_d(NRH)_ under PR conditions were similar (Figs. S2, S3 & S4; Tables S1, S2). Hereafter, we only use *R*_d(Yin)_ for NPR conditions and *R*_d(NRH)_ for PR conditions for further analyses because our study focuses on *R*_d_ and theoretically the Yin method and the NRH method work best to estimate *R*_d_ for NPR and PR conditions, respectively.

### Impact of water regimes on R_d_ and its response to leaf temperature in two wheat genotypes

In EXP2019, *R*_d_ was estimated across two genotypes and two water regimes. As expected, for both NPR and PR conditions, the estimated values of *R*_d_ increased with rising *T*_leaf_ across genotypes and water regimes (Fig. 4), as was widely found in previous studies (Atkin *et al*., 2006; Crous *et al*., 2011; Yin *et al*., 2014; Way *et al*., 2019). The temperature response of *R*_d_ did not vary much between Thésée and Récital but significantly differed between WW and DS conditions. Generally, *R*_d_ was suppressed by drought stress. In agreement with the results of Crous *et al*. (2012), this suppression was slightly or not significant at lower *T*_leaf_ (15 and 20°C), but became increasingly pronounced with increasing *T*_leaf_, reaching up to 76.3% and 71.5% under NPR conditions and up to 74.2% and 64.7% under PR conditions in Thésée and Récital, respectively, for *T*_leaf_ = 40°C (Fig. 4).

**Figure 4.**
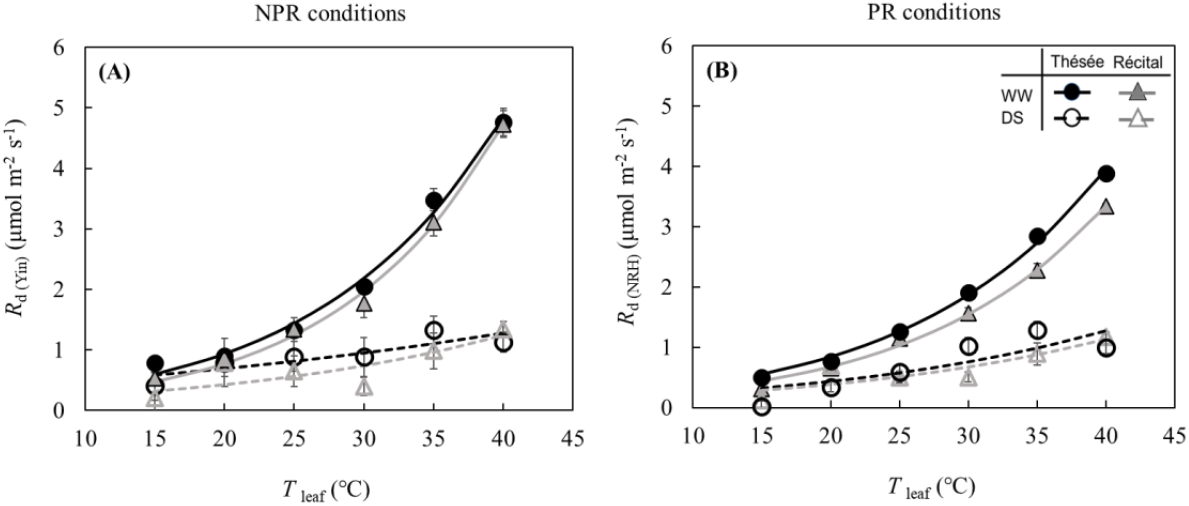
Thermal responses of day respiration estimated by the Yin method (*R*_d(Yin)_) for non-photorespiratory (NPR) conditions (panel A) or by the NRH method (*R*_d(NRH)_ for photorespiratory (PR) conditions (panel B) in two wheat genotypes (Thésée and Récital) under well-watered (WW) and drought-stressed (DS) conditions in EXP2019. The filled points and solid lines represent the WW plants, the open points and dashed lines represent the DS plants. Black symbols and lines refer to Thésée, while gray symbols and lines refer to Récital. Lines are the Arrhenius equation fitted to the data. Error bars indicate standard error of the means (SE) (n = 4).

The temperature response of *R*_d_ was well described by the Arrhenius equation, although the estimated *R*_d_ deviated more under drought conditions as a result of higher variabilities of data among replicated plants. *R*_d_ and its response to *T*_leaf_ showed appreciable acclimation to drought. Under drought stress, *R*_d25_ was apparently reduced, with the reduction ranging from 44.1% to 55.2% across genotypes and between PR and NPR conditions (Fig. 4, Table 1). The values of *E*_Rd_ under well-watered conditions were similar, ranging from 58.87 to 68.64 kJ mol^-1^ in the two genotypes under NPR and PR conditions (Table 1), which agreed with the estimate (64.18 kJ mol^-1^) by Yin *et al*. (2014) for tomato. The estimated *E*_Rd_ was notably affected by water treatments. Under drought conditions the values of *E*_Rd_ decreased to less than 42 kJ mol^-1^ (Table 1), reflecting that *R*_d_ was less sensitive to increasing temperature under water-deficit conditions than in well-watered conditions (Fig. 4).

**Table 1.** Values of modelled leaf day respiration at 25°C (*R*_d25_) and activation energy for leaf day respiration (*E*_Rd_) estimated by the Arrhenius equation for two genotypes of wheat (Thésée and Récital) under well-watered (WW) and drought-stressed (DS) conditions in EXP2019. Data of leaf day respiration used for fitting the Arrhenius equation was estimated by either the Yin method (*R*_d(Yin)_) under non-photorespiratory (NPR) conditions or the NRH method (*R*_d(NRH)_) under photorespiratory (PR) conditions. Standard errors of the estimates are between brackets.

| Condition | Treatment | | $R_{d25}$<br>($\mu\text{mol m}^{-2} \text{s}^{-1}$) | $E_{Rd}$<br>($\text{kJ mol}^{-1}$) | $r^2$ |
| --- | --- | --- | --- | --- | --- |
|  | Genotype | Water regime |  |  |  |
| NPR conditions<br>($R_{d(Yin)}$) | Thésée | WW | 1.45 (0.09) | 62.29 (3.72) | 0.991 |
|  |  | DS | 0.81 (0.08) | 23.53 (7.56) | 0.746 |
|  | Récital | WW | 1.25 (0.06) | 68.64 (3.05) | 0.996 |
|  |  | DS | 0.56 (0.15) | 41.18 (17.74) | 0.615 |
| PR conditions<br>($R_{d(NRH)}$) | Thésée | WW | 1.27 (0.04) | 58.87 (2.00) | 0.997 |
|  |  | DS | 0.58 (0.15) | 40.90 (16.93) | 0.716 |
|  | Récital | WW | 1.04 (0.05) | 60.67 (2.72) | 0.995 |
|  |  | DS | 0.52 (0.10) | 40.94 (12.87) | 0.765 |

### Impact of the combined water and growth temperature regimes on R_d_ and its temperature response

In EXP2020, treatments were designed to investigate the interactive effect of growth temperature and water regime on *R*_d_. The response of *R*_d_ to *T*_leaf_ was described by the Arrhenius equation in each treatment, although it did not fit well the data for DS plants grown at LT (*r*^2^ = 0.329 and 0.395 for NPR and PR conditions, respectively).

Under WW conditions, *R*_d_ estimated at each respective *T*_leaf_ was rather consistently lower in plants grown at HT and MT than in plants grown at LT in both NPR and PR conditions (Fig. 5A,B). This was also reflected in the higher estimation of *R*_d25_ in LT plants under WW conditions (Table 2). However, the estimated *E*_Rd_ for WW plants herein was found to be lowest at LT and highest at HT (Table 2), although the difference between HT and MT was not significant, especially at PR conditions. Our results are contradictory to the lower sensitivity of *R*_d_ to *T*_leaf_ at warmer growth temperature reported previously (Atkin *et al*., 2005 and references therein).

**Figure 5.**
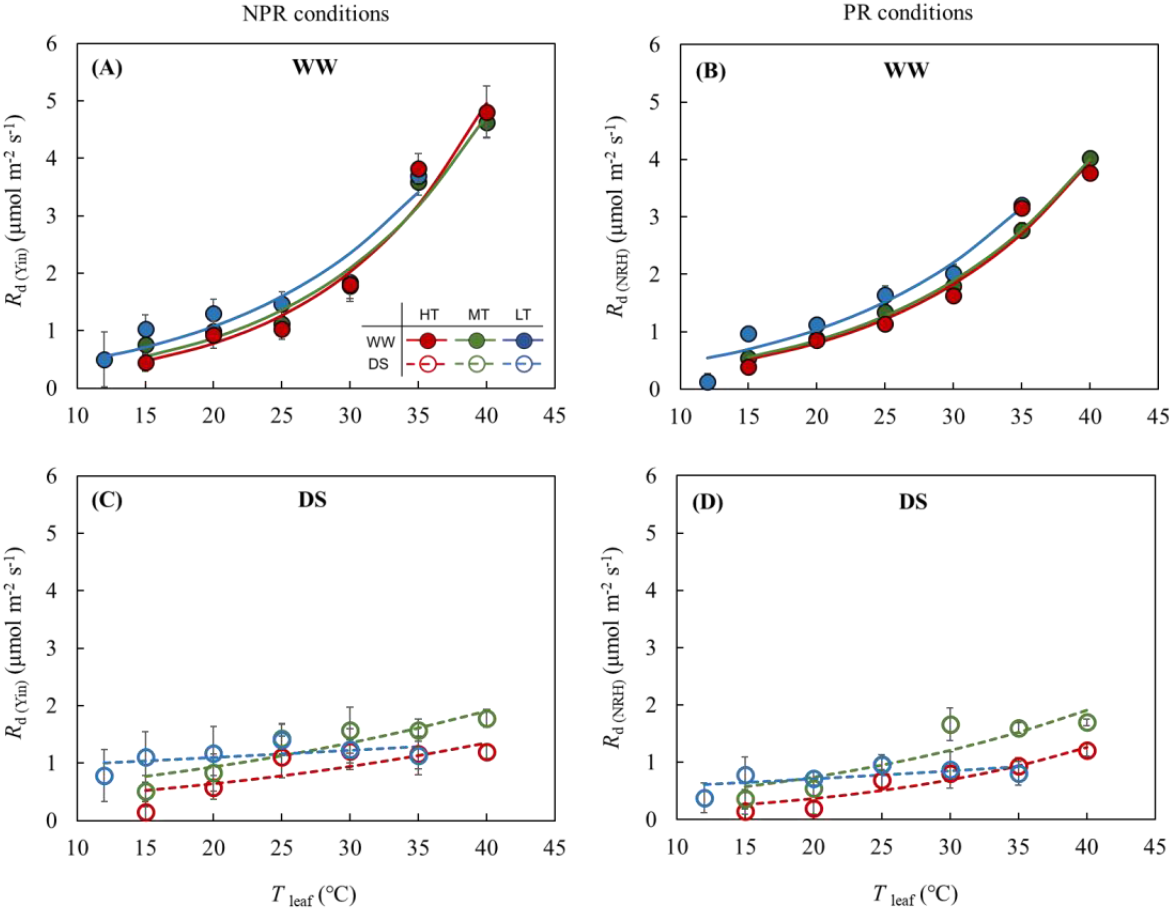
Thermal responses of day respiration estimated by the Yin method (*R*_d(Yin)_) for non-photorespiratory (NPR) conditions (left panels) or by the NRH method (*R*_d(NRH)_ for photorespiratory (PR) conditions (right panels) in winter wheat Thésée grown at three growth temperatures (HT: high temperature, MT: medium temperature, and LT: low temperature) under well-watered (WW; upper panels) and drought-stressed (DS; lower panels) conditions in EXP2020. The filled points and solid lines represent the well-watered plants, the open points and dashed lines represent the drought-stressed plants. Lines are the Arrhenius equation fitted to the data. Error bars indicate standard error of the means (SE) (n=4).

**Table 2.** Values of modelled leaf day respiration at 25°C (*R*_d25_), activation energy for leaf day respiration (*E*_Rd_) estimated by the Arrhenius equation for wheat Thésée grown at three growth temperatures (HT: high temperature, AMT: medium temperature and LT: low temperature) under well-watered (WW) and drought-stressed (DS) conditions in EXP2020. Data of leaf day respiration used for fitting the Arrhenius equation was estimated by either the Yin method (*R*_d(Yin)_) under non-photorespiratory (NPR) conditions or the NRH method (*R*_d(NRH)_) under photorespiratory (PR) conditions. Standard errors of the estimates are between brackets.

| Condition | Treatment | | $R_{d25}$<br>($\mu\text{mol m}^{-2} \text{ s}^{-1}$) | $E_{Rd}$<br>($\text{kJ mol}^{-1}$) | $r^2$ |
| --- | --- | --- | --- | --- | --- |
|  | Genotype | Water regime |  |  |  |
| NPR conditions<br>($R_{d(Yin)}$) | HT | WW | 1.26 (0.15) | 70.69 (7.15) | 0.978 |
|  |  | DS | 0.78 (0.13) | 28.56 (11.86) | 0.672 |
|  | MT | WW | 1.36 (0.16) | 64.58 (7.44) | 0.969 |
|  |  | DS | 1.13 (0.11) | 26.96 (7.27) | 0.816 |
|  | LT | WW | 1.60 (0.18) | 58.00 (10.35) | 0.942 |
|  |  | DS | 1.16 (0.08) | 8.10 (6.09) | 0.329 |
| PR conditions<br>($R_{d(NRH)}$) | HT | WW | 1.22 (0.14) | 60.90 (7.39) | 0.967 |
|  |  | DS | 0.51 (0.08) | 47.10 (9.86) | 0.902 |
|  | MT | WW | 1.27 (0.03) | 59.17 (1.50) | 0.998 |
|  |  | DS | 0.95 (0.14) | 36.09 (10.35) | 0.816 |
|  | LT | WW | 1.52 (0.14) | 55.83 (8.39) | 0.944 |
|  |  | DS | 0.78 (0.07) | 12.93 (8.55) | 0.395 |

Again, drought stress reduced *R*_d_ across various growth environments and between PR and NPR conditions and this reduction was more pronounced at higher *T*_leaf_ (Fig. 5), which led to lower sensitivity to *T*_leaf_, and, thus, lower *E*_Rd_ in DS plants than in WW plants (Table 1). However, this drought-induced reduction of *R*_d_ differed among various growth regimes. DS plants grown at LT maintained relatively higher *R*_d_ at lower *T*_leaf_ (below 25°C) than HT- and MT-grown plants, while at higher *T*_leaf_ (above 30°C) higher *R*_d_ values were observed in DS plants grown at MT as compared with those grown at HT and LT (Fig. 5C,D). This resulted in a downward shift in the temperature response curve and a lower estimated *R*_d25_ (0.78 and 0.51 μmol m^-2^ s^-1^ for NPR and PR, respectively) in DS plants grown at HT; and a nearly horizontal curve with an extremely low estimate of *E*_Rd_ (8.10 and 12.93 kJ mol^-1^ for NPR and PR, respectively) in DS plants grown at LT as compared to those grown at HT and MT (Table 2).

## Discussion

The present study investigated the impacts of long-term drought and growth temperature treatments on the short-term thermal response of *R*_d_ in two winter wheat genotypes. Because little difference in *R*_d_ was observed between the two wheat genotypes under the two contrasting water regimes in EXP2019, only Thésée was used in EXP2020.

### Impact of growth temperature on R_d_ under well-watered conditions

Thermal acclimation of respiration is usually assessed from how changes with growth temperature in the response of respiration rate to measurement *T*_leaf_ sustained over time (Crous *et al*., 2011). In many previous studies, plants acclimated to a sustained warmer climate by reducing their respiration rate at a given measurement temperature (e.g. Crous *et al*., 2011). Moreover, the extent of thermal acclimation of respiration is often lower in pre-existing, fully-expanded leaves that are shifted to a new growth temperature than in leaves that develop under various growth temperatures (Atkin & Tjoelker, 2003). Here, although the plants were subjected to the growth temperature treatment before flag leaves emerged, our results showed that under WW conditions rates of *R*_d_ at any given *T*_leaf_ were nearly identical for plants grown at HT and MT (i.e. no thermal acclimation) regardless of estimation methods or photorespiratory conditions, but *R*_d_ slightly acclimated to the LT with an upward shift of the temperature response curve and a higher *R*_d25_ (Fig. 5A,B, Table 2). This was consistent with findings in previous studies across various *R*_d_ estimation methods and species, indicating that little or only partial thermal acclimation of *R*_d_ is more common (Atkin *et al*., 2006; Way *et al*., 2019).

Plant respiration is sensitive to short-term changes in temperature, while the effect of growth temperature on the thermal sensitivity of respiration varies among species (Atkin *et al*., 2005). Previous works have reported that the sensitivity of respiration to *T*_leaf_ declines with increasing growth temperature, as a result of the acclimation of respiration to a warmer temperature (e.g. Cai *et al*., 2020). In contrast, we found that the estimates of *E*_Rd_ were lower in LT plants than in MT and HT plants under WW conditions (Table 2), implying that plants grown at lower temperatures are likely to be less sensitive to rising *T*_leaf_ than those grown at warmer temperatures. Coincidentally, Crous *et al*. (2017) also observed a higher value of *E*_Rd_ in a fast-growing tree species (*Eucalyptus globulus*) grown in a warmer climate, although the mechanisms behind this remain unclear. The possible explanation of this upward trend of sensitivity to *T*_leaf_ in warm climates could be linked to the higher leaf N content on a mass basis at higher growth temperature (Fig. S5E,F); plant N status is highly associated with factors such as enzyme capacity, substrate supply and respiratory products that may affect the metabolic activities in plant tissues (O’Leary *et al*., 2019 and references therein). Additionally, our data showed that *R*_d25_ positively correlated with leaf N content on area basis (Fig. S5A,B).

### Impact of drought stress on R_d_ and its interaction with growth temperature

Accumulated evidence indicates that the response of respiration and its thermal sensitivity to stressed environments are associated with the alternations in substrate supply, respiratory products demand and respiration/enzyme capacity (Atkin & Tjoelker, 2003; Flexas *et al*., 2005; Atkin & Macherel, 2009). Previous studies suggested that the decreased leaf respiration rate under drought was more likely due to the reduced demand for respiratory products (Ayub *et al*., 2011), while the changed thermal sensitivity was more likely to depend on substrate availability and respiratory enzyme capacity (Atkin & Tjoelker, 2003). The response of leaf respiration to limited water availability seems to be equivocal and elusive as decreased (Ayub *et al*., 2011), unaffected (Gimeno *et al*., 2010) and even increased (Gauthier *et al*., 2014) leaf respiration rate under water deficit have been reported. The various responses could be linked to difference in species used, the severity and duration of soil dehydration and/or other environmental factors, e.g. temperature (Flexas *et al*., 2005).

In our study, inhibition by severe drought on *R*_d_ was slight or not significant under lower *T*_leaf_ across all treatments (Figs. 4,5), consistent with Gauthier *et al*. (2014) who reported that the resilience of leaf respiration in the dark (*R*_dk_) was observed in low to moderate ranges of *T*_leaf_. This finding might explain the unaffected and even slightly increased leaf respiration under drought treatment in some cases where *R*_d_ or *R*_dk_ was measured at a set of common or lower temperature rather than a short-term change of measurement temperature (Gimeno *et al*., 2010; Sperlich *et al*., 2016). At *T*_leaf_ > 20°C, however, our results showed that the inhibition of *R*_d_ by drought was substantial (Figs. 4,5), which was in agreement with previous studies (Haupt-Herting *et al*., 2001; Crous *et al*., 2012; Dahal & Vanlerberghe, 2017). As a result, there was low sensitivity of *R*_d_ to *T*_leaf_ in drought plants (Figs. 4,5). However, several previous studies reported contradictory results: drought may exacerbate *R*_dk_ at moderate or higher temperature (Zagdańska, 1995; Slot *et al*., 2008) and even lead to a ‘respiratory burst’ in an extremely high (> 40°C) short-term measurement temperature range (Gauthier *et al*., 2014), although the underlying mechanisms remain speculative. In some of these studies (e.g. Gauthier *et al*., 2014), plants experienced two phases of drought with a rewatering treatment in between, and the temperature response of leaf respiration was measured at the end of the second period of drought. This means that the increased leaf respiration under drought stress could be due to drought priming. Moreover, in these early studies, drought treatment was applied at seedling or sapling stage (Zagdańska, 1995; Bartoli *et al*., 2005; Gauthier *et al*., 2014), instead of at post-anthesis stage at which assimilated carbohydrates and nitrogenous compounds, those related to substrate supply and respiratory capacity (Tjoelker *et al*., 1999), are being translocated from source (leaves) to sink (grains) (Shao *et al*., 2021), and energy demand for sucrose synthesis and/or phloem loading is in decline (Atkin & Macherel, 2009). The above may explain the difference between their results and our experiments.

Furthermore, our results showed that indeed there was an interactive impact of drought stress and growth temperature on the response of *R*_d_ to *T*_leaf_, with a much lower *E*_Rd_ value in DS plants grown at LT and a lower *R*_d25_ in DS plants grown at HT than in WW plants (Table 2). A low *E*_Rd_ in DS plants grown at LT means a reduced *R*_d_ at higher *T*_leaf_. When *T*_leaf_ was < 25°C, LT treatment appeared to alleviate the negative impact of drought on *R*_d_ rates as compared with HT and MT treatments, whereas this was not the case in the higher *T*_leaf_ range (> 25°C) where DS plants grown at MT exhibited the highest rates of *R*_d_ (Fig. 5C,D). The possible explanation for the mitigated drought effect on LT-grown plants could be linked to the up-regulated alternative oxidase (Searle *et al*., 2011; Dahal & Vanlerberghe, 2017) and mitochondrial uncoupling proteins (Nantes *et al*., 1999; Barreto *et al*., 2017).

### Comparisons of the Kok method, the Yin method and the NRH method

Compared with the Kok method, the Yin method accounts for the additional information from chlorophyll fluorescence data. Our results showed that *R*_d(Kok)_ was lower than *R*_d(Yin)_ under both NPR and PR conditions (Fig. 1A,B), which was in line with previous studies (e.g. Yin *et al*., 2011). The lower estimates of *R*_d(Kok)_ were due to the neglect of a decrease in *Φ*_2_ with increasing light intensity, which occurs even with limiting light levels. Theoretically, the Yin method works well under NPR conditions (Berghuijs *et al*., 2019). It works for PR conditions only if *C*_c_ is maintained constant across light levels, which is technically difficult to achieve in measurements because *g*_m_ is unknown beforehand. Berghuijs *et al*. (2019) pointed out that the linear regression as used in the Yin method will underestimate *R*_d_ if applied to PR conditions; this is also confirmed by our data (Fig. 1D). The theoretical underpinning is that under PR conditions *C*_c_ is not constant but decreases with increasing light intensity, leading to an apparent Kok effect (Farquhar & Busch, 2017; Yin *et al*., 2020). The combined FvCB and *g*_m_ model, Eqn (4), can account for the decline of *C*_c_ with increasing irradiance (Farquhar & Busch, 2017), and thus, in principle, can estimate *R*_d_ for PR conditions. This equation was previously used by Yin & Struik (2009) to estimate *g*_m_, and here we use it to estimate *g*_m_ and *R*_d_ simultaneously, by exploring both gas exchange and chlorophyll fluorescence data across a range of low light intensities after adopting the values of calibration factor *s* (see Eqn (3)) from data measured under NPR conditions. The principle is in analogy to the 2-D modelling of Berghuijs *et al*. (2019) that accounts for the reassimilation of (photo)respired CO_2_, but with the benefit that Eqn (4) is considerably simpler than the 2-D model. It is also in analogy to the procedure of Brooks & Farquhar (1985) that corrects for the decrease in *C*_i_, and of Ayub *et al*. (2011) that further corrects for the decrease in *C*_c_, with increasing irradiance, but with the benefit that the NRH fitting method is easier to implement.

The estimation of *g*_m_ is very sensitive to measurement errors (Yin & Struik, 2009). Here, we failed to estimate *g*_m_ in some cases due to the variation in data from stressed plants (Table S5). Nevertheless, the estimates of *R*_d_ could still be precise as reflected by the low standard error of estimates of *R*_d_. The NRH method has the following advantages. First, this method can estimate *R*_d_ under PR conditions by implicitly considering the variation in *C*_c_ with irradiance as it corrects for the error of the linear regression methods assuming a constant *C*_c_ under PR conditions. Second, this method only requires gas exchange and chlorophyll fluorescence data, but does not require sophisticated and expensive isotopic devices as required by direct *R*_d_ measuring methods (e.g. Haupt-Herting *et al*., 2001; Gong *et al*., 2015) or leaf anatomical data as required by the method of Berghuijs *et al*. (2019). Third, it provides additional estimates for parameters *g*_m_, which could be recognized as indicators of plant physiological processes in response to environmental variables.

Our results showed that light inhibition of *R*_dk_ was higher under PR conditions than under NPR conditions (Fig. 2), consistent with the result of Yin *et al*. (2020) that apparent light inhibition of respiration and thus Kok effect are not obvious under low O_2_ or high CO_2_ conditions or their combinations. Moreover, our data also showed that the underestimation of *R*_d_ by the Kok and Yin methods at photorespiratory conditions may lead to an overestimation of light inhibition of leaf respiration (Fig. 2C,D). Berghuijs *et al*. (2019) suggested that *R*_d_ estimated by the Yin and the Kok methods at NPR conditions cannot represent the real *R*_d_ at PR conditions. Here, our results showed that values of *R*_d_ for NPR conditions were generally higher than those for PR conditions (Fig. 3B,C), which was in agreement with the result of Yin *et al*. (2020) that real light inhibition on respiration increases with increasing amount of photorespiration. Again, the reason for the greater light suppression of *R*_d_ under PR conditions remains to be elucidated.

## Concluding remarks

In contrast to the plethora of studies that have explored the responses of *R*_dk_ to contrasting environments, we assessed the extent to which *R*_d_ of wheat leaves acclimated to drought and growth temperature. We proved a simple method that can estimate *R*_d_ for photorespiratory conditions by using gas exchange and chlorophyll fluorescence data. It was demonstrated that as for non-photorespiratory conditions, *R*_d_ and its temperature response for photorespiratory conditions acclimated more to drought than to growth temperature. Understanding this acclimation of *R*_d_ is needed to support the modelling of *R*_d_, and thus of crop productivity and of carbon cycling in agricultural ecosystems, under future climate change.

## Author contributions

XY and LF conceived the study. LF and PELvdP designed the experiment and conducted the measurements. LF and XY analyzed the data. LF wrote the draft and finalized it with significant input from the other coauthors.

## Notes

### Competing Interest Statement

The authors have declared no competing interest.

