## Supplementary file for "Drought exerts a greater influence than growth temperature on the temperature response of leaf day respiration in wheat (*Triticum aestivum*)"

#### Figures and tables

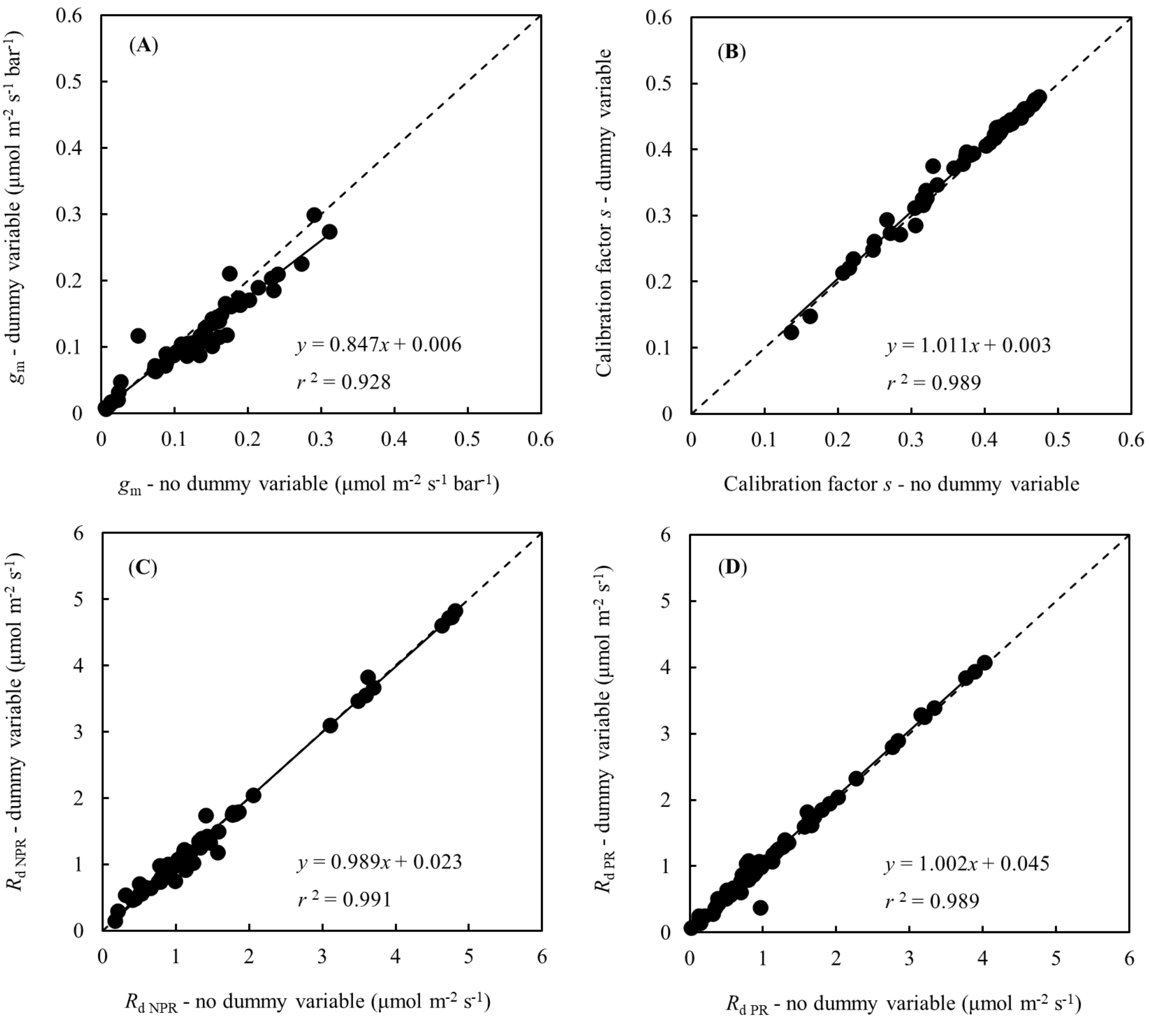


**Figure S1** Correlations between the values estimated by the method with and without dummy variables for parameters mesophyll conductance ($g_m$, panel A), calibrations factor *s* (panel B), leaf day respiration for NPR conditions ($R_{d,NPR}$, panel C) and leaf day respiration for PR conditions ($R_{d,PR}$, panel D) across all treatments. The solid line represents fit to the data of each parameter and the dashed diagonal represents the 1:1 relationship.

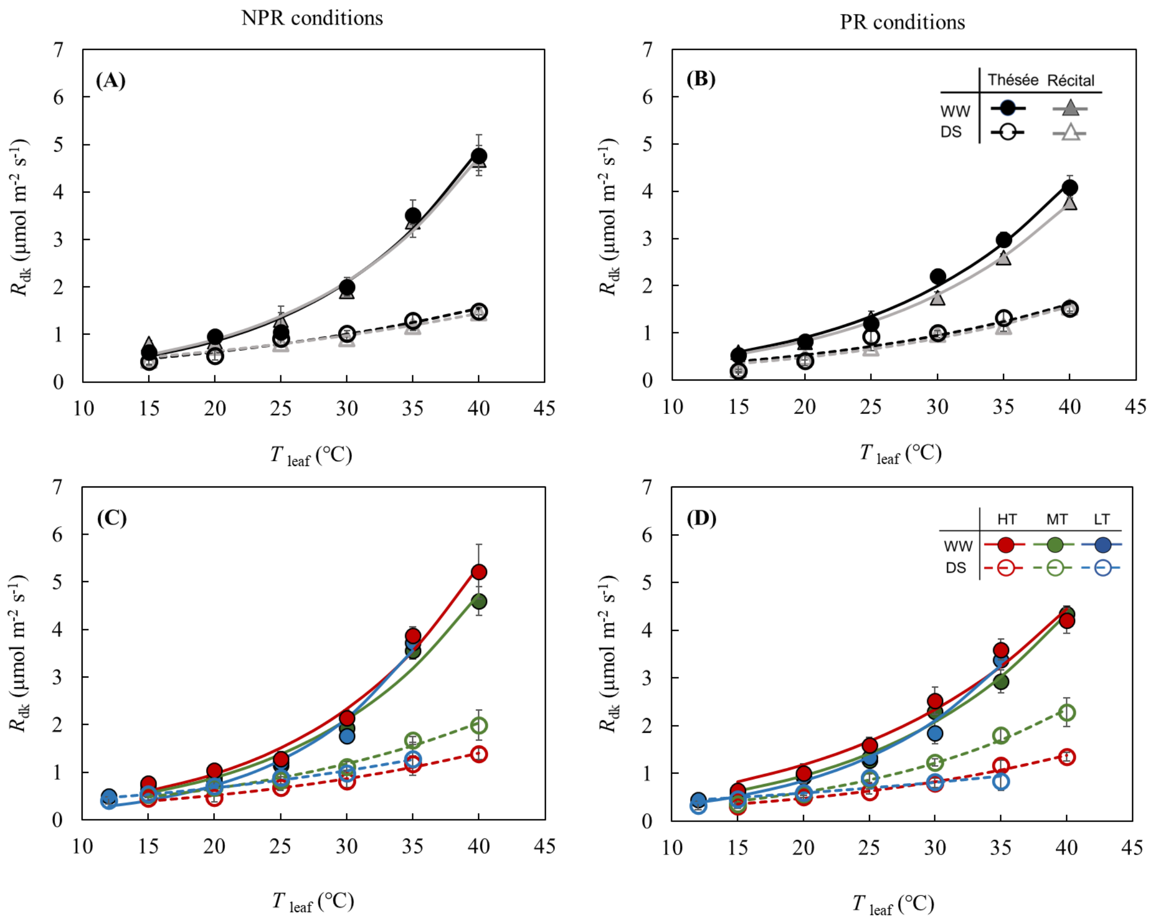


**Figure S2** Thermal responses of respiration in the dark ($R_{dk}$) for non-photorespiratory (NPR; left panels) and photorespiratory (PR; right panels) conditions in two wheat genotypes (Thésée and Récital) under well-watered (WW) and drought-stressed (DS) conditions in EXP2019 (upper panels), and in winter wheat Thésée grown at three growth temperatures (HT: high temperature, MT: medium temperature, and LT: low temperature) under WW and DS conditions in EXP2020 (lower panels). The filled points and solid lines represent the WW plants, the open points and dashed lines represent the DS plants. Lines are the Arrhenius equation fitted to the data. Error bars indicate standard error of the means (SE) (n = 4).

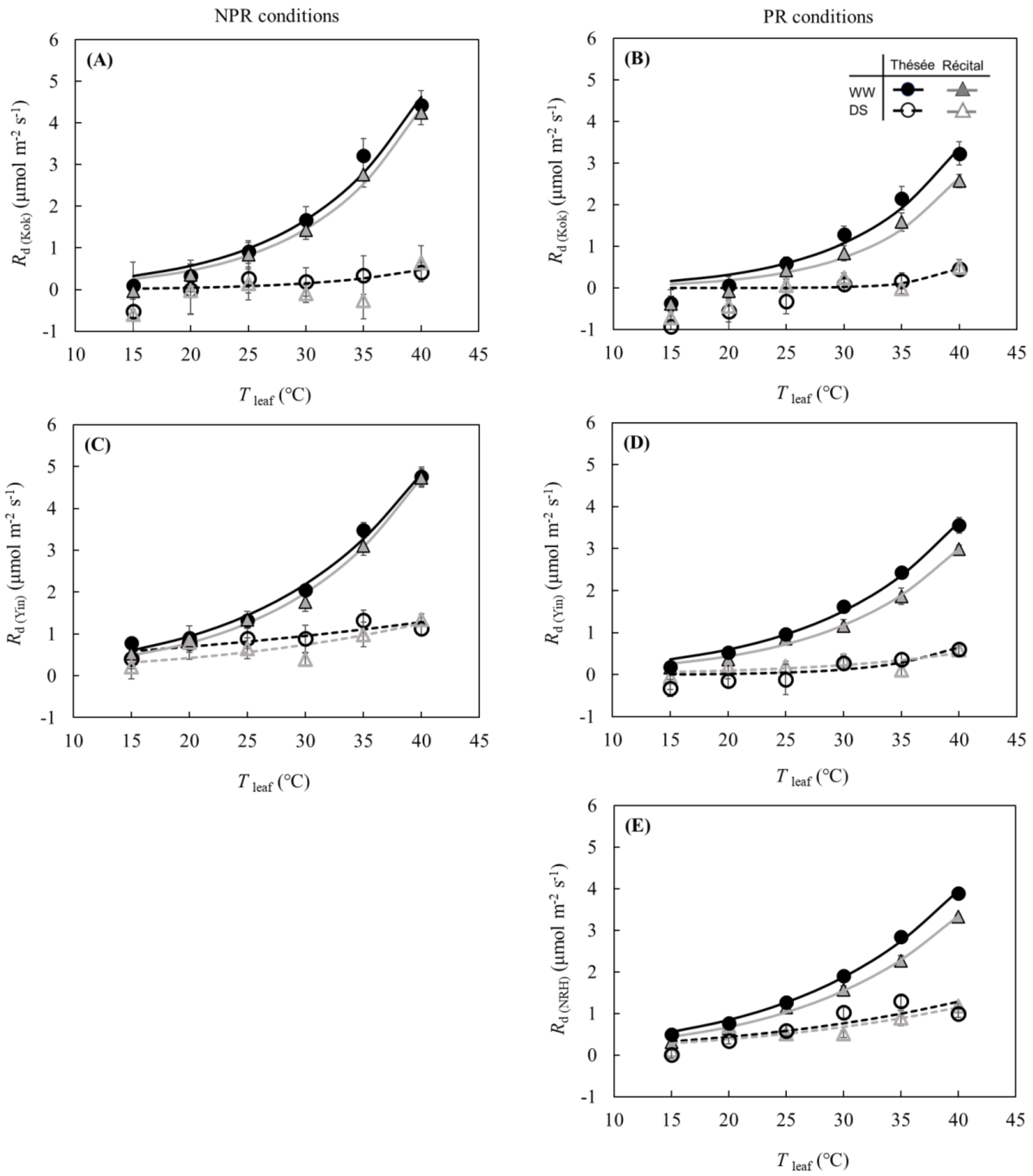


**Figure S3** Thermal responses of leaf day respiration estimated by the Kok method ($R_{d(Kok)}$) and by the Yin method ($R_{d(Yin)}$) for non-photorespiratory (NPR; panels A and C) or thermal responses of $R_{d(Kok)}$, $R_{d(Yin)}$ and leaf day respiration estimated by the NRH method ($R_{d(NRH)}$) for photorespiratory (PR; panels B, D and E) conditions in two wheat genotypes (Thésée and Récital) under well-watered (WW) and drought-stressed (DS) conditions in EXP2019. The filled points and solid lines represent the WW plants, the open points and dashed lines represent the DS plants. Lines are the Arrhenius equation fitted to the data. In panels A and B, thermal response of genotype Récital grown at DS conditions could not be fitted by the Arrhenius equation due to the high variability of data. Error bars indicate standard error of the means (SE) (n = 4).

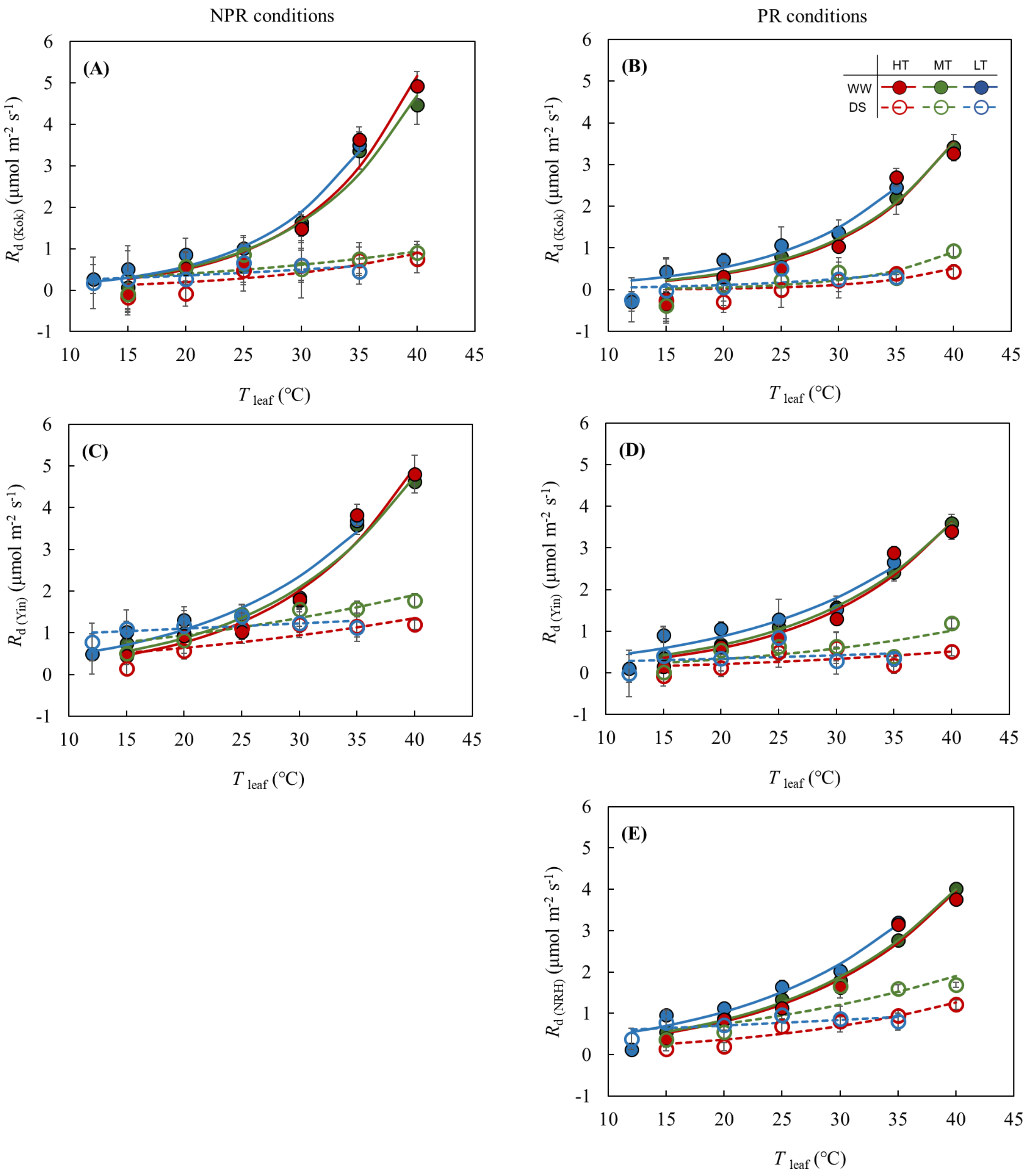


**Figure S4** Thermal responses of day respiration estimated by the Kok method ($R_{d(Kok)}$) and by the Yin method ($R_{d(Yin)}$) for non-photorespiratory (NPR; panels A and C) or thermal responses of $R_{d(Kok)}$, $R_{d(Yin)}$ and leaf day respiration estimated by the NRH method ($R_{d(NRH)}$) for photorespiratory (PR; panels B, D and E) conditions in winter wheat Thésée grown at three growth temperatures (HT: high temperature, MT: medium temperature, and LT: low temperature) under WW and DS conditions in EXP2020. The filled points and solid lines represent the WW plants, the open points and dashed lines represent the DS plants. Lines are the Arrhenius equation fitted to the data. Error bars indicate standard error of the means (SE) (n = 4).

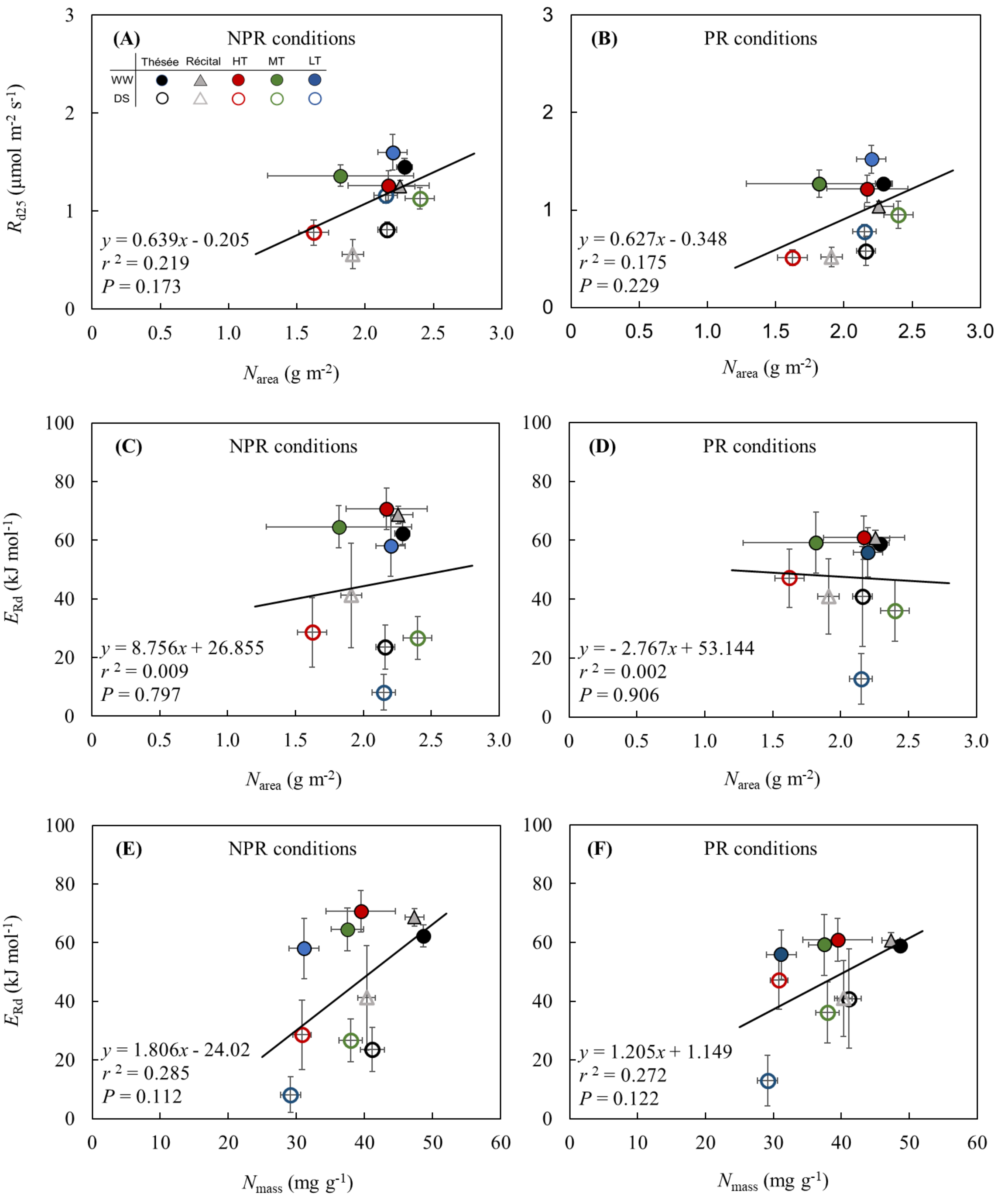


**Figure S5** Correlations between values of modelled leaf day respiration at 25 °C ($R_{d25}$; panels A and B) or values of activation energy for leaf day respiration ($E_{Rd}$; panels C and D) and leaf nitrogen content on area basis ($N_{area}$), and correlations between $E_{Rd}$ and leaf nitrogen content on mass basis ($N_{mass}$; panels E and F), under non-photorespiratory (NPR; left panels) and photorespiratory (PR; right panels) conditions. Error bars indicate standard error of the estimates or stand error of the means. The solid line represents fit to the data. $R_{d25}$ and $E_{Rd}$ in this figure were estimated by the $R_d$ data derived from the Yin method under NPR conditions and from the NRH method under PR conditions.

**Table S1** Values of parameters estimated at 25 °C ($X_{25}$) and activation energy ($E_x$) estimated by the Arrhenius equation for two genotypes of wheat (Thésée and Récital) under well-watered (WW) and drought-stressed (DS) conditions in EXP2019. Data used for fitting the Arrhenius equation was presented in Figure S3. Standard errors of the estimates are between brackets.

| | Treatment | | | | |
|---|---|---|---|---|---|
| | Genotype | Water regime | $X_{25}$ (μmol m$^{-2}$ s$^{-1}$) | $E_x$ (kJ mol$^{-1}$) | $r^2$ |
| $R_{dk}$ (NPR) | Thésée | WW | 1.35 (0.12) | 66.36 (5.55) | 0.991 |
| | | DS | 0.80 (0.04) | 33.76 (3.42) | 0.969 |
| | Récital | WW | 1.39 (0.10) | 63.33 (4.36) | 0.989 |
| | | DS | 0.80 (0.03) | 30.61 (2.65) | 0.975 |
| $R_{dk}$ (PR) | Thésée | WW | 1.35 (0.08) | 58.46 (3.59) | 0.991 |
| | | DS | 0.72 (0.09) | 42.22 (8.02) | 0.913 |
| | Récital | WW | 1.24 (0.03) | 57.15 (1.52) | 0.998 |
| | | DS | 0.65 (0.04) | 45.36 (4.24) | 0.977 |
| $R_{d(Kok)}$ (NPR) | Thésée | WW | 0.98 (0.14) | 80.16 (8.50) | 0.960 |
| | | DS | 0.08 (0.15) | 90.53 (103.70) | 0.494 |
| | Récital | WW | 0.82 (0.10) | 86.27 (6.78) | 0.991 |
| | | DS | -* | -* | -* |
| $R_{d(Yin)}$ (NPR) | Thésée | WW | 1.45 (0.09) | 62.29 (3.72) | 0.991 |
| | | DS | 0.81 (0.08) | 23.53 (7.56) | 0.746 |
| | Récital | WW | 1.25 (0.06) | 68.64 (3.05) | 0.996 |
| | | DS | 0.56 (0.15) | 41.18 (17.74) | 0.615 |
| $R_{d(Kok)}$ (PR) | Thésée | WW | 0.60 (0.17) | 89.30 (16.56) | 0.960 |
| | | DS | 0.01 (0.08) | 226.60 (747.40) | 0.524 |
| | Récital | WW | 0.38 (0.14) | 101.20 (20.84) | 0.958 |
| | | DS | -* | -* | -* |
| $R_{d(Yin)}$ (PR) | Thésée | WW | 0.96 (0.07) | 68.75 (4.18) | 0.993 |
| | | DS | 0.05 (0.09) | 129.10 (89.57) | 0.804 |
| | Récital | WW | 0.72 (0.05) | 73.67 (3.96) | 0.994 |
| | | DS | 0.14 (0.08) | 65.82 (35.18) | 0.599 |
| $R_{d(NRH)}$ (PR) | Thésée | WW | 1.27 (0.04) | 58.87 (2.00) | 0.997 |
| | | DS | 0.58 (0.15) | 40.90 (16.93) | 0.716 |
| | Récital | WW | 1.04 (0.05) | 60.67 (2.72) | 0.995 |
| | | DS | 0.52 (0.10) | 40.94 (12.87) | 0.765 |

* Thermal response could not be fitted by the Arrhenius equation due to the high variability of data.

NPR stands for non-photorespiratory conditions; PR stands for photorespiratory conditions.

**Table S2** Values of parameters estimated at 25 °C ($X_{25}$) and activation energy ($E_x$) estimated by the Arrhenius equation for wheat Thésée grown at three growth temperatures (HT: high temperature, AMT: medium temperature and LT: low temperature) under well-watered (WW) and drought-stressed (DS) conditions in EXP2020. Data used for fitting the Arrhenius equation was presented in Figure S4. Standard errors of the estimates are between brackets.

| | Treatment | | | | |
|---|---|---|---|---|---|
| | Genotype | Water regime | $X_{25}$ (µmol m$^{-2}$ s$^{-1}$) | $E_x$ (kJ mol$^{-1}$) | $r^2$ |
| $R_{dk}$ (NPR) | HT | WW | 1.51 (0.13) | 65.07 (5.10) | 0.986 |
| | | DS | 0.67 (0.03) | 38.18 (2.98) | 0.981 |
| | MT | WW | 1.38 (0.14) | 63.73 (6.20) | 0.978 |
| | | DS | 0.88 (0.04) | 42.97 (2.96) | 0.986 |
| | LT | WW | 1.26 (0.15) | 79.45 (10.32) | 0.967 |
| | | DS | 0.84 (0.02) | 31.95 (2.53) | 0.979 |
| $R_{dk}$ (PR) | HT | WW | 1.67 (0.14) | 50.65 (5.32) | 0.973 |
| | | DS | 0.63 (0.03) | 40.63 (2.89) | 0.985 |
| | MT | WW | 57.99 (3.17) | 57.99 (3.17) | 0.992 |
| | | DS | 51.45 (2.67) | 51.45 (2.67) | 0.993 |
| | LT | WW | 67.64 (5.73) | 67.64 (5.73) | 0.983 |
| | | DS | 22.89 (8.09) | 22.89 (8.09) | 0.709 |
| $R_{d(Kok)}$ (NPR) | HT | WW | 0.94 (0.21) | 88.42 (13.13) | 0.966 |
| | | DS | 0.28 (0.13) | 59.64 (29.99) | 0.711 |
| | MT | WW | 0.95 (0.19) | 82.51 (11.85) | 0.966 |
| | | DS | 0.50 (0.14) | 33.13 (20.14) | 0.498 |
| | LT | WW | 1.06 (0.15) | 87.91 (12.48) | 0.964 |
| | | DS | 0.42 (0.07) | 25.24 (15.94) | 0.454 |
| $R_{d(Yin)}$ (NPR) | HT | WW | 1.26 (0.15) | 70.69 (7.15) | 0.978 |
| | | DS | 0.78 (0.13) | 28.56 (11.86) | 0.672 |
| | MT | WW | 1.36 (0.16) | 64.58 (7.44) | 0.969 |
| | | DS | 1.13 (0.11) | 26.96 (7.27) | 0.816 |
| | LT | WW | 1.60 (0.18) | 58.00 (10.35) | 0.942 |
| | | DS | 1.16 (0.08) | 8.10 (6.09) | 0.329 |
| $R_{d(Kok)}$ (PR) | HT | WW | 0.67 (0.23) | 85.92 (19.81) | 0.930 |
| | | DS | 0.06 (0.10) | 114.40 (103.10) | 0.708 |
| | MT | WW | 0.71 (0.13) | 82.82 (10.56) | 0.977 |
| | | DS | 107.60 (56.20) | 107.60 (56.20) | 0.804 |
| | LT | WW | 76.24 (16.24) | 76.24 (16.24) | 0.921 |
| | | DS | 63.05 (67.18) | 63.05 (67.18) | 0.413 |
| $R_{d(Yin)}$ (PR) | HT | WW | 0.95 (0.17) | 68.76 (10.89) | 0.951 |
| | | DS | 0.26 (0.12) | 34.64 (32.15) | 0.329 |
| | MT | WW | 1.04 (0.03) | 64.14 (1.59) | 0.999 |
| | | DS | 0.44 (0.14) | 42.99 (21.07) | 0.581 |
| | LT | WW | 1.27 (0.14) | 53.29 (10.68) | 0.900 |
| | | DS | 0.39 (0.12) | 15.43 (29.68) | 0.089 |
| $R_{d(NRH)}$ (PR) | HT | WW | 1.22 (0.14) | 60.90 (7.39) | 0.967 |
| | | DS | 0.51 (0.08) | 47.10 (9.86) | 0.902 |
| | MT | WW | 1.27 (0.03) | 59.17 (1.50) | 0.998 |
| | | DS | 0.95 (0.14) | 36.09 (10.35) | 0.816 |
| | LT | WW | 1.52 (0.14) | 55.83 (8.39) | 0.944 |
| | | DS | 0.78 (0.07) | 12.93 (8.55) | 0.395 |

NPR stands for non-photorespiratory conditions; PR stands for photorespiratory conditions.

**Table S3** Values of calibration factor *s* (standard error of the estimate between brackets) estimated by the Yin method under non-photorespiratory (NPR) conditions across all treatments and leaf temperatures in leaves of wheat Thésée and Récital. WW: well-watered; DS: drought-stressed; HT: high temperature; MT: medium temperature; LT: low temperature.

| | **Treatment** | | **Leaf temperature** | | | | | | |
|---|---|---|---|---|---|---|---|---|---|
| | **Genotype** | **Water regime** | **12 °C** | **15 °C** | **20 °C** | **25 °C** | **30 °C** | **35 °C** | **40 °C** |
| EXP 2019 | Thésée | WW | NA | 0.458 (0.007) | 0.457 (0.007) | 0.469 (0.006) | 0.447 (0.006) | 0.445 (0.010) | 0.432 (0.014) |
| | | DS | NA | 0.385 (0.016) | 0.402 (0.015) | 0.377 (0.015) | 0.318 (0.020) | 0.271 (0.017) | 0.206 (0.016) |
| | Récital | WW | NA | 0.445 (0.006) | 0.474 (0.004) | 0.465 (0.011) | 0.466 (0.012) | 0.468 (0.012) | 0.453 (0.014) |
| | | DS | NA | 0.335 (0.024) | 0.422 (0.024) | 0.320 (0.017) | 0.274 (0.011) | 0.284 (0.025) | 0.249 (0.011) |
| | **Growth temperature** | **Water regime** | | | | | | | |
| EXP 2020 | HT | WW | NA | 0.432 (0.008) | 0.406 (0.013) | 0.435 (0.009) | 0.446 (0.013) | 0.428 (0.013) | 0.422 (0.024) |
| | | DS | NA | 0.313 (0.020) | 0.315 (0.014) | 0.358 (0.022) | 0.326 (0.012) | 0.229 (0.021) | 0.242 (0.010) |
| | MT | WW | NA | 0.437 (0.010) | 0.451 (0.010) | 0.448 (0.015) | 0.419 (0.013) | 0.438 (0.011) | 0.440 (0.016) |
| | | DS | NA | 0.413 (0.022) | 0.374 (0.025) | 0.383 (0.016) | 0.305 (0.025) | 0.305 (0.012) | 0.215 (0.013) |
| | LT | WW | 0.416 (0.027) | 0.466 (0.014) | 0.454 (0.012) | 0.449 (0.011) | 0.414 (0.006) | 0.442 (0.008) | NA |
| | | DS | 0.375 (0.035) | 0.428 (0.033) | 0.370 (0.028) | 0.329 (0.017) | 0.161 (0.024) | 0.136 (0.023) | NA |

NA not applicable

**Table S4** Values of calibration factor *s'* (standard error of the estimate between brackets) estimated by the Yin method under photorespiratory (PR) conditions across all treatments and leaf temperatures in leaves of wheat Thésée and Récital. WW: well-watered; DS: drought-stressed; HT: high temperature; MT: medium temperature; LT: low temperature.

| | **Treatment** | | **Leaf temperature** | | | | | | |
|---|---|---|---|---|---|---|---|---|---|
| | **Genotype** | **Water regime** | **12 °C** | **15 °C** | **20 °C** | **25 °C** | **30 °C** | **35 °C** | **40 °C** |
| EXP 2019 | Thésée | WW | NA | 0.338 (0.003) | 0.334 (0.005) | 0.322 (0.006) | 0.300 (0.003) | 0.271 (0.005) | 0.256 (0.011) |
| | | DS | NA | 0.251 (0.012) | 0.252 (0.010) | 0.178 (0.019) | 0.132 (0.009) | 0073 (0.008) | 0.077 (0.012) |
| | Récital | WW | NA | 0.352 (0.004) | 0.344 (0.004) | 0.326 (0.006) | 0.301 (0.007) | 0.285 (0.010) | 0.263 (0.005) |
| | | DS | NA | 0.234 (0.021) | 0.274 (0.017) | 0.179 (0.008) | 0.138 (0.007) | 0.080 (0.009) | 0.084 (0.008) |
| | **Growth temperature** | **Water regime** | | | | | | | |
| EXP 2020 | HT | WW | NA | 0.323 (0.008) | 0.298 (0.011) | 0.301 (0.006) | 0.292 (0.005) | 0.267 (0.008) | 0244 (0.010) |
| | | DS | NA | 0.219 (0.007) | 0.227 (0.014) | 0.226 (0.020) | 0.169 (0.020) | 0.056 (0.013) | 0.071 (0.008) |
| | MT | WW | NA | 0.335 (0.009) | 0.333 (0.006) | 0.314 (0.005) | 0.279 (0.008) | 0.273 (0.010) | 0.252 (0.012) |
| | | DS | NA | 0.271 (0.024) | 0.264 (0.025) | 0.208 (0.019) | 0.109 (0.018) | 0.074 (0.003) | 0.092 (0.008) |
| | LT | WW | 0.320 (0.019) | 0.357 (0.010) | 0.352 (0.008) | 0.269 (0.023) | 0.239 (0.014) | 0.260 (0.008) | NA |
| | | DS | 0.224 (0.039) | 0.263 (0.049) | 0.206 (0.017) | 0.182 (0.016) | 0.048 (0.016) | 0.041 (0.009) | NA |

NA not applicable

**Table S5** Values of $g_m$ (mol m$^{-2}$ s$^{-1}$ bar$^{-1}$, standard error of the estimate between brackets) estimated by the NRH-A method under photorespiratory (PR) conditions across all treatments and leaf temperatures in leaves of wheat Thésée and Récital. WW: well-watered; DS: drought-stressed; HT: high temperature; MT: medium temperature; LT: low temperature.

| | **Treatment** | | **Leaf temperature** | | | | | | |
|---|---|---|---|---|---|---|---|---|---|
| | **Genotype** | **Water regime** | **12 °C** | **15 °C** | **20 °C** | **25 °C** | **30 °C** | **35 °C** | **40 °C** |
| EXP 2019 | Thésée | WW | NA | 0.109 (0.010) | 0.158 (0.029) | 0.164 (0.040) | 0.202 (0.040) | 0.109 (0.019) | 0.099 (0.065) |
| | | DS | NA | 0.089 (0.018) | 0.188 (0.061) | 0.131 (0.040) | 0.088 (0.034) | 0.013 (0.003) | 0.006 (0.002) |
| | Récital | WW | NA | 0.175 (0.034) | 0.152 (0.019) | 0.177 (0.050) | 0.142 (0.036) | 0.118 (0.039) | 0.076 (0.014) |
| | | DS | NA | 0.160 (0.173) | 0.214 (0.139) | 0.142 (0.063) | -# | 0.023 (0.007) | 0.022 (0.006) |
| | **Growth temperature** | **Water regime** | | | | | | | |
| EXP 2020 | HT | WW | NA | 0.119 (0.024) | 0.160 (0.092) | 0.171 (0.039) | 0.169 (0.041) | 0.151 (0.069) | 0.074 (0.026) |
| | | DS | NA | 0.073 (0.031) | 0.117 (0.075) | 0.147 (0.109) | -# | 0.011 (0.003) | 0.008 (0.002) |
| | MT | WW | NA | 0.232 (0.155) | 0.273 (0.119) | 0.311 (0.116) | 0.235 (0.118) | 0.189 (0.120) | 0.072 (0.032) |
| | | DS | NA | 0.125 (0.076) | 0.134 (0.131) | 0.135 (0.073) | 0.050 (0.033) | 0.026 (0.005) | 0.017 (0.008) |
| | LT | WW | 0.087 (0.026) | 0.241 (0.133) | 0.320 (0.183) | 0.290 (0.182) | 0.151 (0.066) | 0.102 (0.029) | NA |
| | | DS | 0.088 (0.054) | -# | -# | -# | 0.006 (0.003) | 0.005 (0.003) | NA |

NA not applicable

### $g_m$ failed to be estimated together with $R_d$ due to the high variability of data under extreme conditions.
